# GBP1 promotes killing of multidrug-resistant *Acinetobacter baumannii* and promotes host protection via interferon signalling and inflammasome activation

**DOI:** 10.1101/2022.10.24.513490

**Authors:** Fei-Ju Li, Lora Starrs, Anukriti Mathur, Daniel Enosi Tuipulotu, Si Ming Man, Gaetan Burgio

## Abstract

Multidrug resistant (MDR) *Acinetobacter baumannii* are of major concern worldwide due to their resistance to last resort carbapenem and polymyxin antibiotics. To develop an effective treatment strategy, it is critical to better understand how an *A. baumannii* MDR bacterium interacts with its mammalian host. Pattern-recognition receptors sense microbes, and activate the inflammasome pathway, leading to pro-inflammatory cytokine production and programmed cell death. Here, we found that MDR *A. baumannii* activate the NLRP3 inflammasome complex predominantly via the non-canonical caspase-11-dependent pathway. We show that caspase-11-deficient mice are protected from a virulent MDR *A. baumannii* strain by maintaining a balance between protective and deleterious inflammation via IL-1. Caspase-11-deficient mice also compromise between effector cell recruitment, phagocytosis, and programmed cell death in the lung during infection. Importantly, we found that cytosolic immunity - mediated by guanylate-binding protein 1 (GBP1) and type I interferon signalling - orchestrates caspase-11-dependent inflammasome activation. This exerts a bactericidal activity against carbapenem- and colistin-resistant, lipooligosaccharide (LOS)- deficient bacteria. Together, our results suggest that developing therapeutic strategies targeting GBP1 might pave the way as a host-directed therapy to overcome multidrug resistance in *A. baumannii* infection.

## Introduction

*Acinetobacter baumannii* is a Gram-negative bacterium and has emerged as one of the most prevalent causative agents of nosocomial infections around the world ^1^, frequently leading to urinary tract infections, intensive care unit (ICU)-acquired pneumonia and septicemia ^2 3^. In the USA alone, ICU-acquired *A. baumannii* pneumonia presents a considerable disease burden, being encountered in encountered in 5-10 % of patients receiving mechanical ventilation ^4^ and resulting in a high fatality rate due to septicemia ^5^. It is classified by the World Health Organization (WHO) as a member of the ESKAPE pathogens (*Enterococcus faecium, Staphylococcus aureus, Klebsiella pneumoniae, Acinetobacter baumannii, Pseudomonas aeruginosa* and *Enterobacter spp*.). Due to carbapenem and colistin antibiotic resistance, *A. baumannii* is listed amongst the strains that are critical for new therapeutic strategies by WHO^1^. Unfortunately, beyond combination therapies, there are currently no efficacious treatments against multidrug resistant (MDR) and extensively drug resistant (XDR) *A. baumannii* bacteria ^6^. Targeting the host instead of the pathogen, solely or as a combination therapy, could potentially lead to novel avenues in overcoming – and potentially further circumventing - MDR/XDR resistance. To identify potential host targets, a comprehensive characterisation of the mechanisms of the host innate response to the MDR/XDR *A. baumannii* bacteria is required.

Once *A. baumannii* invades the host, stimuli such as pathogen-associated molecular patterns (PAMPs), dead cells or irritants (danger-associated molecular patterns, DAMPs) are detected and the host mounts a protective inflammatory response via inflammasomes ^7^. PAMPs and DAMPs are sensed by cytosolic inflammasome sensors such as Absent in Melanoma 2 (AIM2), NOD-like receptors (NLRs such as NLRC4 and NLRP3 ^8^). These inflammasome sensors recruit the inflammasome adaptor protein apoptosis-associated speck-like protein containing a caspase activation and recruitment domain (ASC, also known as PYCARD) upon activation ^9^. The resultant inflammasome complex activates caspase-1, which is required to induce cleavage of the pro-inflammatory cytokines pro-interleukin-1β (pro-IL-1β) and pro-IL-18, as well as the pro-pyroptotic factor, Gasdermin D (GSDMD) ^10,11^, which along with NINJ1 ^12^ drives an inflammatory programmed cell death known as pyroptosis ^13,14^. Previous work reported that the NLRP3 inflammasome via the ‘canonical’ caspase-1 pathway contributes to host defence against *A. baumannii* isolates in an intranasal infection model ^15^. However, NLRP3 only partially protected the host against *A. baumannii* bacteria strains suggesting the existence of additional protective mechanisms against the bacteria.

A ‘non-canonical’ NLRP3 inflammasome activation pathway has been described that is dependent on caspase-11 in mice ^11^ and caspase-4 and -5 in humans ^16^. This pathway is essential to defend against cytosolic pathogens via interferon (IFN) signalling ^17^. IFNs activate multiple host cell death pathways ^18^ suggesting a potential protective role for cytosolic immunity against MDR *A. baumannii* infection. Cytosolic immunity is an important host innate defence mediated by bacterial lipopolysaccharide (LPS) and interferon-inducible proteins, such as the bactericidal GTPase guanylate-binding-proteins (GBPs) in the cytoplasm ^19^. However, the role and the mechanism of non-canonical inflammasome activation and these cytosolic defence systems in response to MDR *A. baumannii* infection are unknown.

Here we performed a comprehensive characterisation of the non-canonical inflammasome and cytosolic immune response to a virulent *A. baumannii* MDR strain (ATCC BAA-1605) resistant to carbapenem and a lipooligosaccharide (LOS)-deficient clinical strain resistant to polymyxin (ATCC 19606 strains) ^20 21^. We firstly found both canonical and caspase-11 non-canonical NLRP3 pathways were activated in response to systemic and severe infection. We discovered that caspase-11-deficiency conferred a full protective effect against the infection and mediated the balance between protective versus detrimental hyper-inflammation via IL-1β, but not IL-18. Intriguingly, we found that the host utilises type I IFN signalling to mediate the caspase-11 non-canonical NLRP3 inflammasome activation. This then activates the interferon-inducible GBP1 bactericidal activity against carbapenem and LOS-deficient polymyxin resistant *A. baumannii* strains. Together these findings identified the requirement of caspase-11 and cytosolic immunity in mediating the inflammatory response and in exerting a bactericidal activity against MDR and virulent *A. baumannii* strains.

## Results

### The multidrug resistant strain *A. baumannii* ATCC BAA-1605 activates caspase-1 and caspase-11 inflammasomes and induces programmed cell death

Previous work has reported that *A. baumannii* membrane proteins elicit an innate immune response by inducing the expression of pro-inflammatory cytokines IL-1β and IL-18 ^22^. Indeed, we noted elevated IL-1β and IL-18 cytokine secretion levels in primary wild-type (WT) bone marrow derived macrophages (BMDMs) infected with the multidrug resistant (MDR) virulent *A. baumannii* ATCC BAA-1605 strain (hereafter named *A. baumannii* 1605). To identify the inflammasome sensors responsible for the recognition of *A. baumannii*, we inoculated WT BMDMs with *A. baumannii* 1605 and measured *Nlrc4, Aim2. Nlrp3* and *Caspase-11* gene expression levels in cell lysates at 6 and 12 hours post infection. We found a sustained high expression in *Nlrp3* and *Casp11* transcript levels over time. These data suggest that *Nlrp3* and *Casp11* are possible sensors for *A. baumannii* 1605 (**Fig. 1b**). To confirm this finding, we inoculated *Nlrp3*^*-/-*^, *Asc*^*-/-*^, *Casp11*^*-/*-^ and *Casp1/11*^*-/-*^ BMDMs with *A. baumannii* 1605. We noted a strong reduction to an abolition of IL-1β secretion in all knockouts, and a significant decrease in IL-18 in *Casp11*^*-/*-^ and *Casp1/11*^*-/-*^ BMDMs. In contrast, the release of TNFα for BMDMs from all WT and knockout BMDMs was maintained (**Fig. 1a**). These data suggest an activation of NLRP3 inflammasome via caspase-1 and/or caspase-11 pathways. Indeed, we found an activation of both caspase-1 and caspase-11 in WT BMDMs via immunoblotting (**Fig. 1c**). Activation of caspase-1 and caspase-11 cleaves the N-terminal end of Gasdermin D (GSDMD), resulting in a mature GSDMD, which forms membrane pores leading to pyroptosis ^10^. We observed GSDMD cleavage in WT BMDMs (**Fig. 1c**), increased cell death (**Fig. 1d**) and LDH release (**Fig. 1e**). Mature GSDMD formation was almost abolished in *Casp11*^*-/-*^ and *Casp1/11*^*-/-*^ BMDMs (**Fig. 1c**). Programmed cell death (**Fig. 1d**), but not LDH release, was reduced in *Gsdmd*^*-/-*^ BMDMs compared to WT BMDMs (**Fig. 1e**). Collectively these data indicate that *A. baumannii* 1605 strain induces both activation of caspase-11 leading to pro-inflammatory cytokine secretion, Gasdermin D proteolytic cleavage, programmed cell death and activation of the NLRP3-Caspase-1 inflammasome.

**Figure 1.**
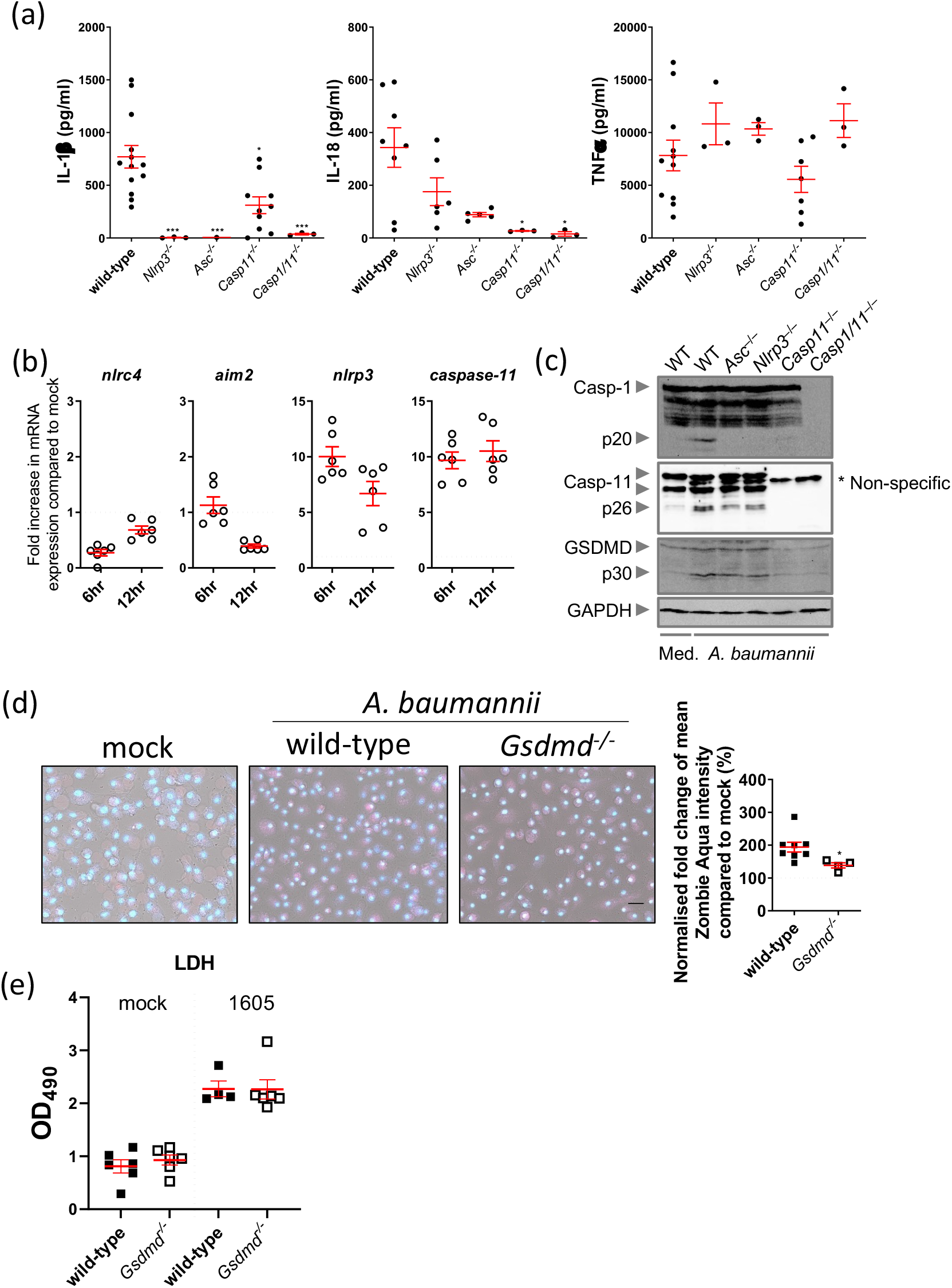
*A. baumannii 1605* bacteria induce inflammasome activation. **(a)** Cytokine levels IL-1β, IL-18 and TNFα in supernatants from WT and mutant mouse BMDMs 12 hours post infection (MOI = 10), n = 3-13 biological replicates, each dot represents one replicate. **(b)** Transcript levels of inflammasome sensor genes *Nlrc4, Aim2, Nlrp3* and *Caspase-11* produced in wild-type mouse BMDMs 6 hours post infection (MOI = 10) and measured by quantitative PCR, n = 6, mean ± SEM. **(c)** Western blots on activated caspases and GSDMD 16 hours post infection (MOI = 10). **(d)** Immunofluorescence imaging of BMDMs cell death post 24 hours of infection (MOI = 10). Scale bar 30 um. (e) LDH release by mouse BMDMs 12 hours post-infection, n=6, mean ± SEM. *, P < 0.05, **, P < 0.01, ***, P < 0.001 compared to wild-type. mean ± SEM. Non-parametric t-test was used to compare differences between groups.

### Virulent MDR *A. baumannii* 1605 infection predominantly activates caspase-11-NLRP3 inflammasome

To confirm the preferential inflammasome activation pathway by virulent MDR *A. baumannii* 1605, we inoculated *Casp1*^*-/-*^, *Casp11*^*-/*-^, *Casp1/11*^*-/*-^, *Asc*^*-/-*^ and *Nlrp3*^*-/-*^ mice with *A. baumannii* 1605 intra-peritoneally at 2×10^7^ CFU/mouse and measured survival, bacterial burden, and plasma IL-18 or IL-1β levels. Although *Casp1*^*-/-*^ (**Fig. 2a**) and *Casp1/11*^*-/*-^ (**Supp. Fig. 2a**) exhibited a similar survival rate (45% survival rate) and similar bacterial burden to WT, both *Nlrp3*^*-/-*^ and *Asc*^*-/-*^ mice were protected (60% survival rate) against *A. baumannii* 1605 (**Fig. 2a**). Remarkably, we found 80% survival to the infection **(Fig 2a)**, a significant reduction of the bacteria burden (**Fig. 2b, 2c** and **Supp. Fig. 2b**) and reduction of the plasma pro-inflammatory cytokine secretion (**Fig. 2d** and **Supp. 2c**) in *Casp11*^-/-^ mice. Interestingly, bacterial burden and pro-inflammatory cytokine secretion were also significant lower in *Asc*^*-/-*^ mice compared to WT mice, but these levels were not as low as in *Casp11*^-/-^ mice in most organs. We next examined the role of GSDMD-mediated cell death by infecting *Gsdmd*^*-/-*^ mice. *A. baumannii* 1605 bacteria inoculation in *Gsdmd*^*-/-*^ mice resulted in a 60% survival rate when compared to WT mice (**Supp. Fig. 3c and d**). While the bacterial burden in the *Gsdmd*^*-/-*^ mice did not differ from WT mice, we however found a 10-fold reduction in plasmatic IL-18 in *Gsdmd*^*-/-*^ mice (**Supp. Fig. 3b**). We similarly found a 10-fold reduction in IL-18 level in the lysates of infected *Gsdmd*^*-/-*^, and a marginal decrease in IL-1β and TNFα secretion level (**Supp. Fig. 3a**). Taken together this genetic validation experiment suggests *A. baumannii* 1605 activates caspase-11 resulting in the activation of the NLRP3/ASC inflammasome, GSDMD, and the induction of programmed cell death. Additionally, these data confirmed that while NLRP3/ASC or caspase-11 deficiency alone conferred mouse survival, deficiency in both caspase-1 and caspase-11 did not protect against the infection.

**Figure 2.**
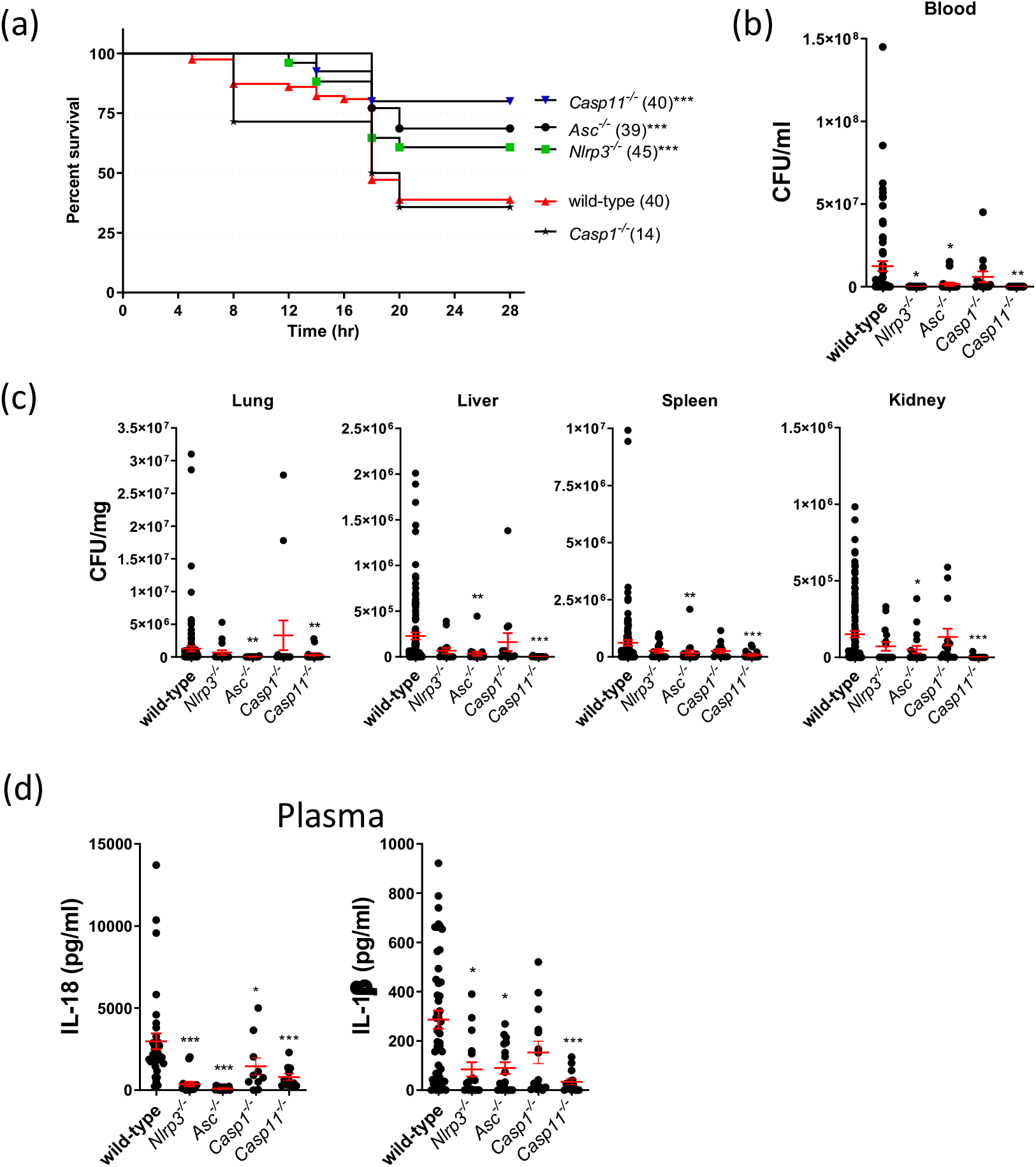
Inflammasome deficient mice confer resistance to *A. baumannii* 1605 infection. **(a)** Survival rate of WT, *Nlrp3*^*-/-*^, *Asc*^*-/-*^, *Caspase1*^*-/-*^ and *Caspase11*^*-/-*^ mice 28 hours post infection (i.p. 2×10^7^ CFU/mouse). **(b)** The bacteremia and **(c)** bacteria dissemination to different organs (lung, liver, spleen or kidney) at 10-20 hours post infection were quantified by serial dilution on trypticase soy broth and CFU counting. **(d)** Cytokine levels IL-1β and IL-18 in plasma 12 hours post infection. Data were collected from at least three independent experiments, numbers of mice (n) are indicated in parentheses, *, P < 0.05, **, P < 0.01, ***, P < 0.001 compared to wild-type. mean ± SEM. Kaplan-Meier estimate was used to compare mice survival rates. Non-parametric t-test was used to compare differences between groups.

**Figure 3.**
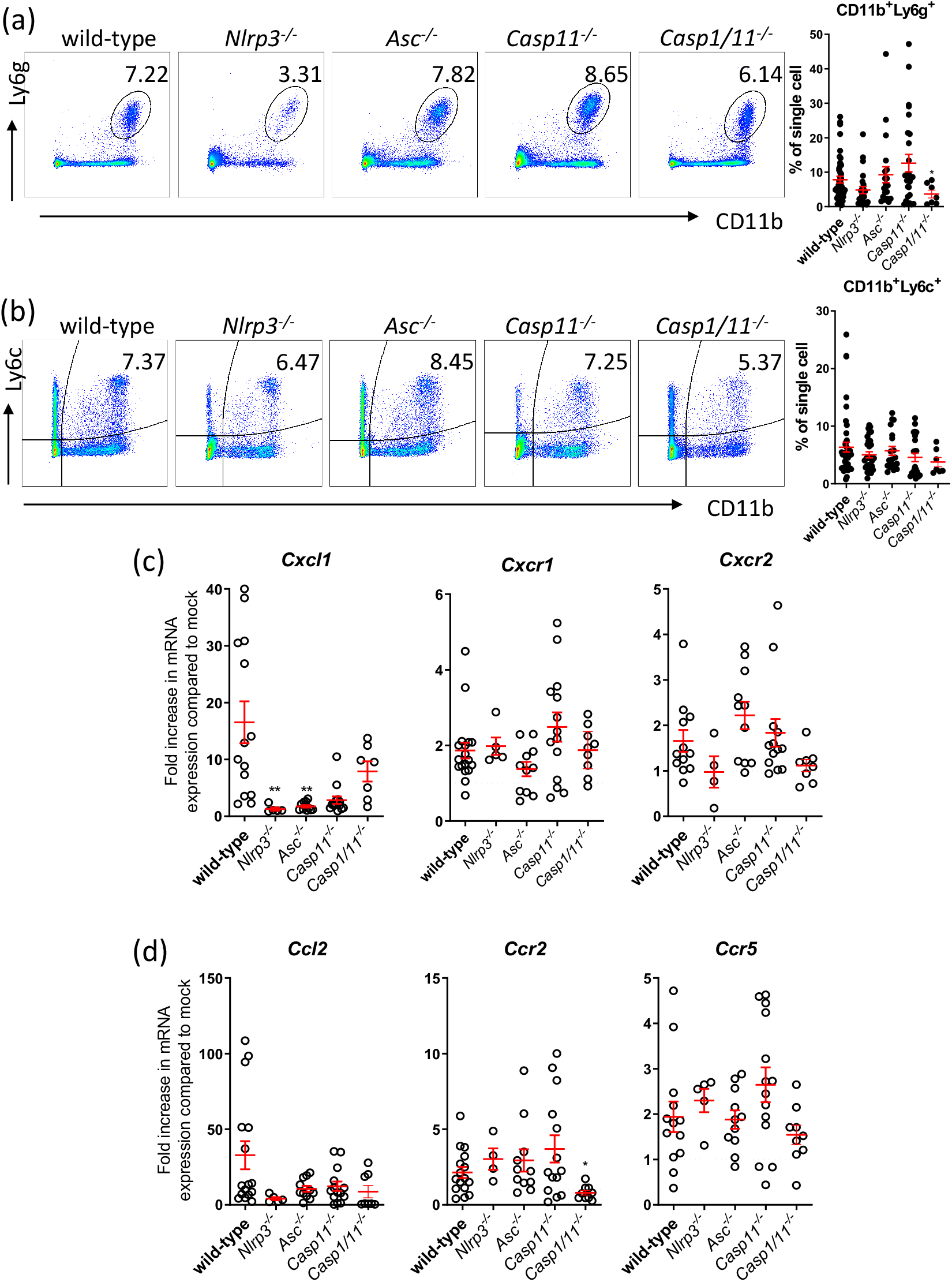
Absence of NLRP3 signalling decreases neutrophil recruitment to the lungs. Flow cytometry quantification of **(a)** neutrophils (CD11b^+^Ly6g^+^) and **(b)** inflammatory monocytes (CD11b^+^Ly6c^+^Ly6g^-^), in the mice lung 14-20 hours infection (i.p. 2×10^7^ CFU/mouse), n = 20. qPCR quantification of induction of **(c)** neutrophil chemokine and chemokine receptors or **(d)** inflammatory monocyte chemokine and receptors. Data were collected from at least three independent experiments, n = 4-14 for each group, each data point represents a replicate. *, P < 0.05, mean ± SEM. Non-parametric t-test was used to compare differences between groups.

### Reduced programmed cell death in the lung partly underlies host resistance to *A. baumannii* 1605 infection

We next sought to determine how *Casp11*^*-/-*^, *Nlrp3*^*-/-*^ and *Asc*^*-/-*^ mice were protected against the infection, whereas *Casp1*^*-/-*^ and *Casp1/11*^*-/-*^ were not. Previous works have reported that during lung infection, depletion of neutrophils resulted in an acute lethal infection ^23^, and macrophage depletion lead to an increased tissue bacterial burden ^24^. We therefore postulated that increased neutrophils (CD11b^+^Ly6g^+^) and inflammatory monocytes (CD11b^+^Ly6c^+^) recruitment and/or increased clearance of infected cells in the target tissues, such as the lung, were responsible for enhanced resistance to *A. baumannii* 1605. While we identified - by flow cytometry - an increase of CD11b^+^Ly6g^+^ and CD11b^+^Ly6c^+^ populations in the lungs of WT mice from mock infected to six-hour post infection, and again from six-hour to 20-hour post inoculation (**Supp. Fig. 4a**), we observed no difference in the percentage of CD11b^+^Ly6g^+^ and CD11b^+^Ly6c^+^ in the lungs between WT, *Asc*^*-/-*^, *Casp1/11*^*-/-*^ and *Casp11*^*-/-*^ mice (**Fig. 3a** and **3b**). Interestingly, we noted a reduction in CD11b^+^Ly6g^+^ but not CD11b^+^Ly6c^+^ cell population in the lungs of the *Nlrp3*^*-/-*^ mice (**Fig. 3a**) and a reduction in chemokine *Cxcl1* expression level, markers of neutrophils recruitment in *Nlrp3*^*-/-*^ and *Asc*^*-/-*^ mice (**Fig. 3c**). We observed the expression levels of the neutrophil chemokine receptors *Cxcr1* and *Cxcr2* remained similar in the lungs between the WT and the four strains of knockout mice (**Fig. 3c**). There was a significant decrease in the expression of the inflammatory monocyte marker, *Ccr2*, in *Casp1/11*^*-/-*^ mice, consistent with low inflammation observed in *Casp1/11*^*-/-*^ mice **(Suppl Fig 2d)**. However, we observed no difference between the WT and knockout mice in the expression of two other markers of inflammatory monocyte recruitment *Ccl2* and *Ccr5* **(Fig. 3d**). We also observed no difference in the pathology of the lungs (**Supp. Fig. 4 b-d**). Taken together, these findings suggest that increased neutrophil and inflammatory monocyte recruitment are unlikely to play a major role in the survival of *Asc*^*-/-*^ and *Casp11*^*-/-*^ mice. In contrast, the reduction in neutrophil recruitment in *Nlrp3*^*-/-*^ mice might have contributed to preventing the progression of lethal *A. baumannii* 1605-mediated inflammation.

**Figure 4.**
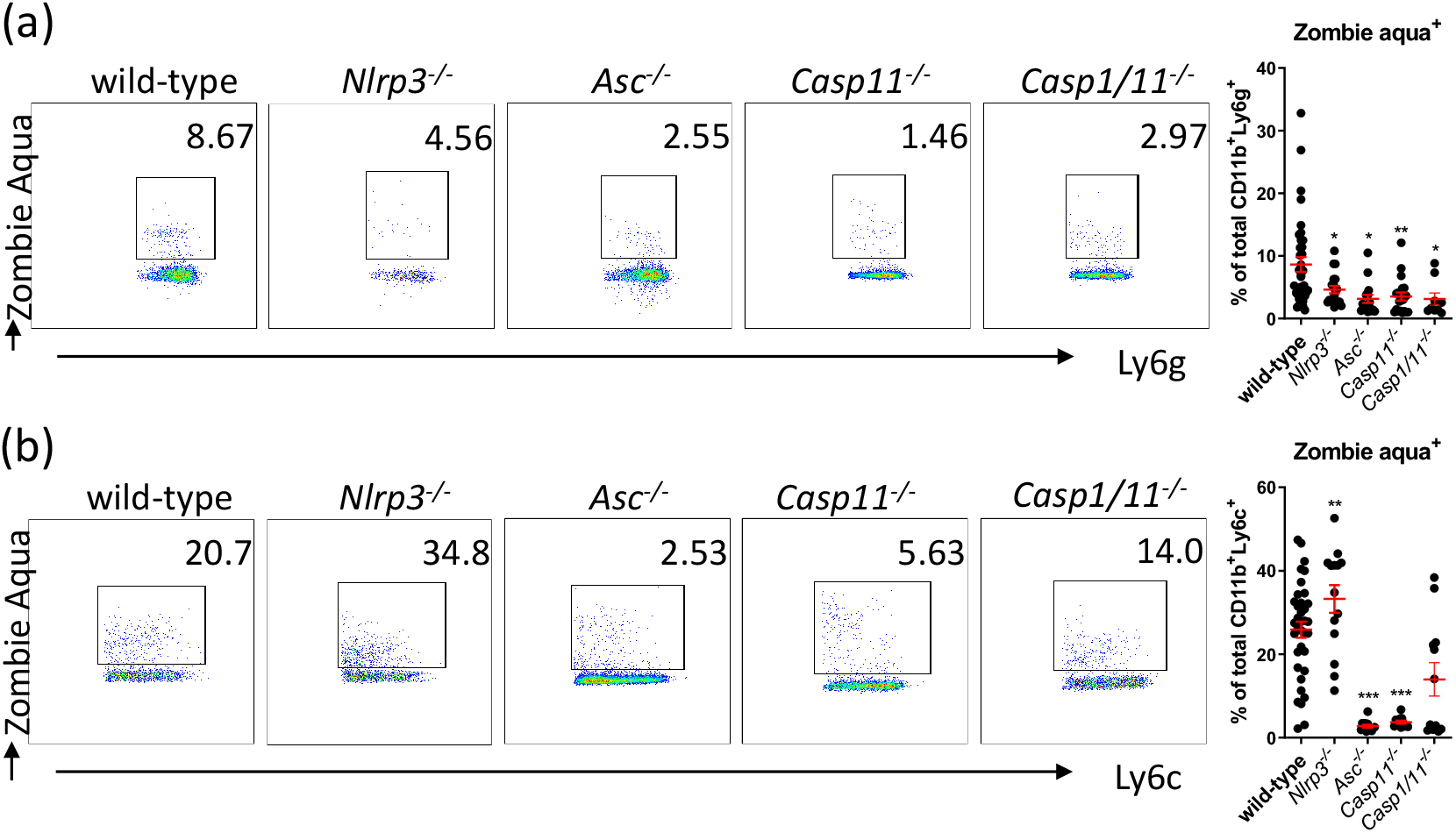
Absence of inflammasome signalling decreases effector cell death. Flow cytometry quantification of **(a)** neutrophils (CD11b^+^Ly6g^+^) cell death (Zombie Aqua^+^) and **(b)** inflammatory monocytes (CD11b^+^Ly6c^+^Ly6g^-^) cell death, in mice lung 14-20 hours infection (i.p. 2×10^7^ CFU/mouse). *, P < 0.05, **, P < 0.01, ***, P < 0.001, mean ± SEM, n = 20. Non-parametric t-test was used to compare differences between groups.

We next postulated that reduction in effector cell death might have instead protected *Nlrp3*^*-/-*^, *Asc*^*-/-*^, and *Casp11*^*-/-*^ mice against severe inflammation. We quantified the percentage of neutrophils undergoing programmed cell death in the lungs of these mice using Zombie aqua dye. Interestingly, all four knock-out strains showed a significant decrease in the number of dead or dying neutrophils (**Fig 4a**). Remarkably, while we found no difference in Zombie aqua-positive inflammatory monocytes between WT and *Casp1/11*^*-/-*^ mice, we observed a significant increase in the number of dead or dying inflammatory monocytes in *Nlrp3*^*-/-*^ mice compared to WT. The significantly lower Zombie aqua-positive cells for *Asc*^*-/-*^ and *Casp11*^*-/-*^ mice suggests a reduced inflammatory monocyte cell death in these strains than in WT mice **(Fig. 4b)**.

These data collectively suggest that in *Nlrp3*^-/-^, *Asc*^*-/-*^ and *Casp11*^*-/-*^ mice, there may be a tight balance between recruitment of neutrophils and/or inflammatory monocytes to the lungs, and that the ability of these effectors to phagocytose and undergo programmed cell death may contribute to the improved bacterial clearance and decreased host mortality.

### Inflammasome-dependent IL-1 mediates the balance of protective versus deleterious inflammation in response to *A baumannii* 1605 severe sepsis

Pro-inflammatory cytokines are key mediators for protecting the host against infection ^25^. However, a dysregulation in pro-inflammatory cytokine production leads to excessive inflammation and severe sepsis ^26^. Conversely, a lack of pro-inflammatory cytokine production results in increased bacterial burden and severe bacteremia ^26^. We speculated that dysregulation of inflammasome-dependant pro-inflammatory cytokine (IL-1 and IL-18) secretion might have been responsible for severe sepsis and lethality in mice infected with MDR *A. baumannii* 1605. We assessed the role of inflammasome-dependent interleukin (IL) signalling pathway by blocking IL-18 ^27^ and IL-1 receptor (IL-1R) ^28^ in *II-18*^*-/-*^, *Il-1r1*^*-/-*^ knockout mice, and both pathways in *Il-1r1/18*^*-/-*^ knockout mice. We inoculated *A. baumannii* 1605 bacteria into these knockout mice and monitored the infection up to 28 hours post inoculation (**Fig. 5a**). We observed that the survival rate for *Il-18*^*-/-*^ and *Il-1r1/18*^*-/-*^ were similar or lower than WT mice. We, however, found a striking 90% survival rate for *Il-1r1*^*-/-*^ mice (**Fig. 5a**) and a 4 to 10-fold bacterial load reduction in *Il-1r1*^*-/-*^ and *Il-18*^*-/-*^ within the lung and liver. Interestingly, we found an increased bacterial burden in *Il-1r1/18*^*-/-*^ mice (**Fig. 5b-c**). These findings, suggest that inflammasome-dependent IL-1, and not IL-18, mediates a tight balance between beneficial and detrimental inflammation in the host in response to virulent MDR *A. baumannii* 1605 infection.

**Figure 5.**
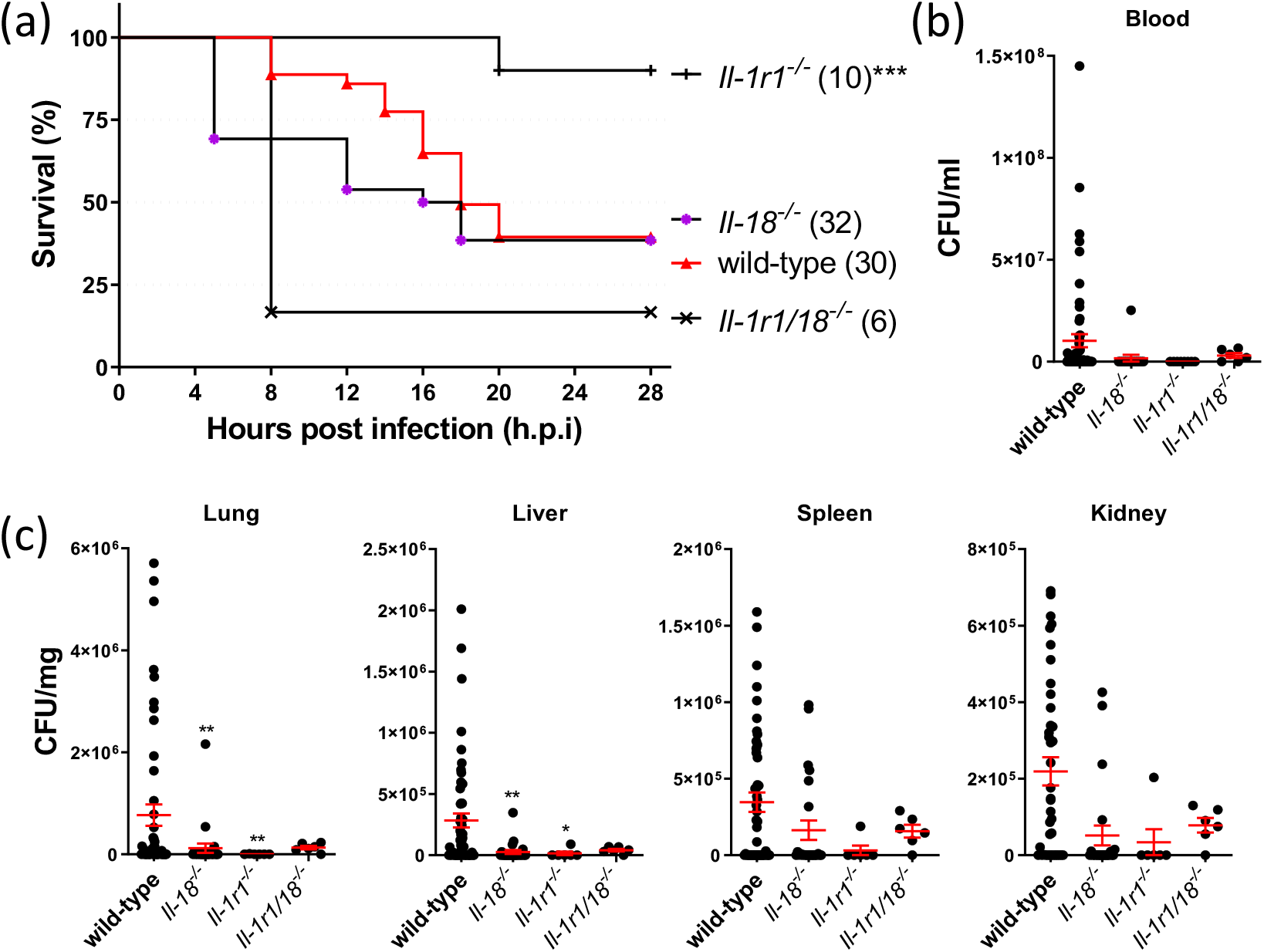
Deleterious inflammation drives acute lethality in *A. baumannii*-infected mice. Mouse **(a)** survival rate, **(b)** the level of bacteremia and **(c)** bacteria dissemination to different organs 16-20 hours post *A. baumannii* 1605 infection (i.p. 2×10^7^ CFU/mouse). Data were collected from at least three independent experiments, number of mice (n) as indicated in parentheses, *, P < 0.05, **, P < 0.01, ***, P < 0.001 compared to wild-type. mean ± SEM, numbers of mice (n) are indicated in parentheses. Kaplan-Meier estimate was used to compare mice survival rates. Non-parametric t-test was used to compare differences between groups.

### Type I IFN is required for host protection and bactericidal activity against MDR *A. baumannii* 1605

A previous study and our group have reported that activation of caspase-11 in *A. baumannii* infection is dependent on type I IFN signalling ^29^. We postulated that type I IFN priming is critical in *A. baumannii* 1605-induced caspase-11 non-canonical pathway activation resulting in detrimental inflammation. We inoculated the *Ifnar*^*-/-*^ BMDMs with *A. baumannii* 1605 and measured *Nlrp3* and *Caspase-11* transcript levels and pro-inflammatory cytokine levels up to 12 hours post infection. We observed a reduction in *Caspase-11* transcript levels while *Nlrp3* transcript levels were unchanged (**Fig. 6b**). Additionally, we noted a lower to abolished pro-inflammatory cytokine transcripts and secretion (**Fig. 6a** and **Supp. Fig. 5**) in *Ifnar*^*-/-*^ infected mice suggesting that IFN primes caspase-11 activation, 12 hours post *A. baumannii* 1605 infection. Next, we inoculated WT and *Ifnar*^*-/-*^ mice intraperitoneally with *A. baumannii* 1605 bacteria at 2×10^7^ CFU/mouse. We assessed survival, bacterial load and plasmatic pro-inflammatory cytokine secretion. Remarkably, we found almost full protection of the *Ifnar*^*-/-*^ mice with 80% survival rate 28 hours post inoculation (**Fig. 6c**). Interestingly, we observed a significant reduction in bacterial load (blood, lung, liver and kidneys; **Fig. 6d-e**) and quasi abolition of the pro-inflammatory plasma cytokine levels in *Ifnar*^*-/-*^ mice (**Fig. 6f**). These results suggest there is a protective role if type I IFN signalling is abolished during infection with MDR *A. baumannii* 1605. We next sought to determine the mechanism/s of resistance of the *Ifnar*^*-/-*^ mice. We analysed CD11b^+^Ly6g^+^ and CD11b^+^Ly6c^+^ populations, chemokine receptor levels and Zombie aqua-positive cells in the lungs of *Ifnar*^*-/-*^ and WT mice, 16-20 hours post infection. We found a significant reduction in CD11b^+^Ly6g^+^ population (**Supp. Fig. 6a**) but not CD11b^+^Ly6c^+^ (**Supp. Fig. 7a**) and a substantial reduction in Zombie aqua-positive cells in CD11b^+^Ly6g^+^ and CD11b^+^Ly6c^+^ cell types in *Ifnar*^*-/-*^ compared to WT mice (**Supp. Fig. 6c** and **7c**). Interestingly, we found no difference in chemokine levels (**Supp. Fig. 6b** and **7b**). Together these findings confirm that an IFN-dependent caspase-11 response is required following MDR *A. baumannii* 1605 infection to mediate the release of pro-inflammatory cytokines and encourage the persistence of effector cells in target organs.

**Figure 6.**
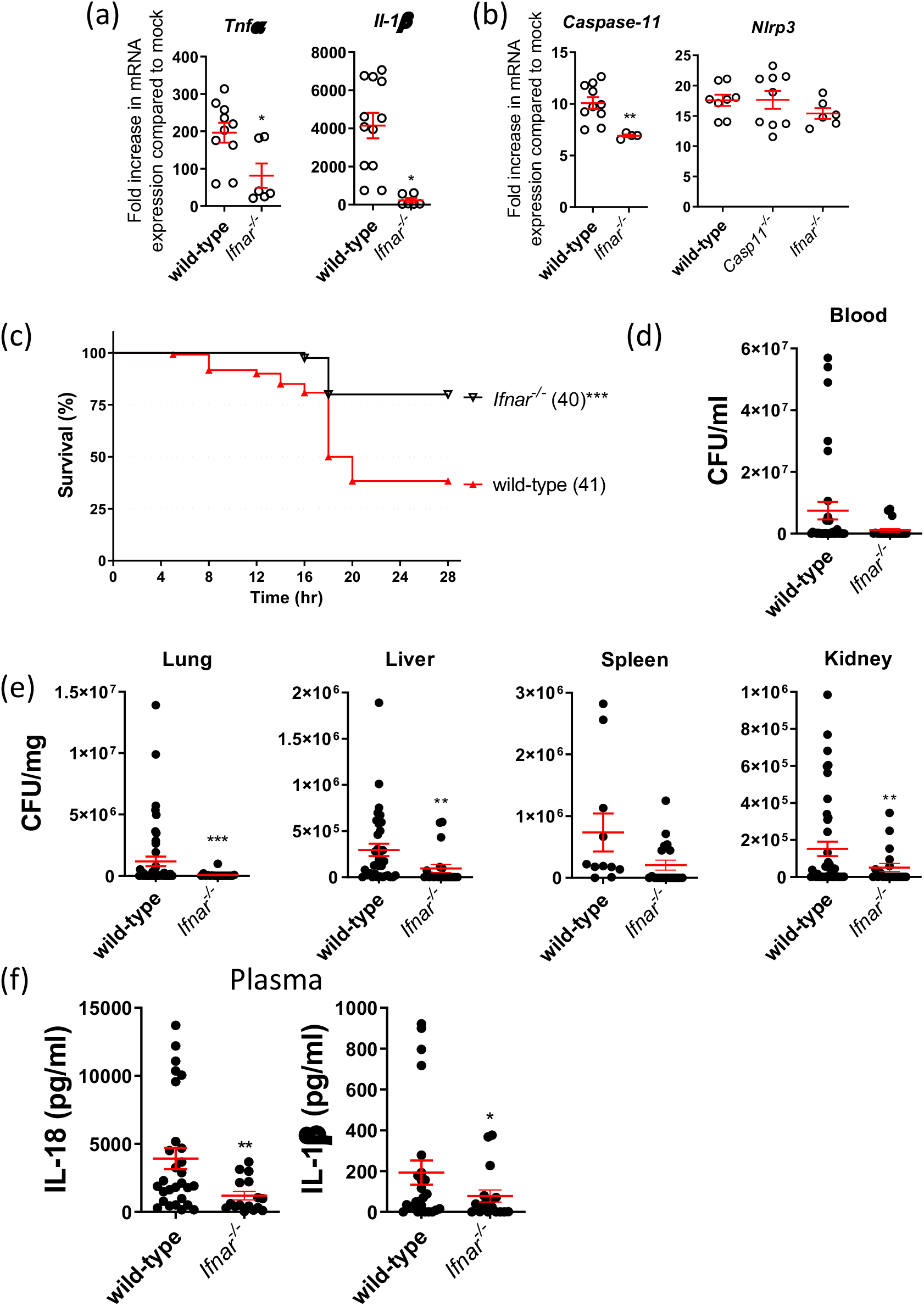
*A. baumannii* induces type I IFN-dependent inflammasome activation. Transcript levels measured by quantitative PCR of **(a)** inflammatory cytokines and **(b)** inflammasome produced by mouse BMDM after 6 hours of infection (MOI = 10), n=6-12, each data point represents a replicate. **(c)** Mice survival rate, **(d)** the level of bacteraemia, **(e)** bacteria dissemination to different organs, and **(f)** plasma cytokine levels 16-20 hours post *A. baumannii* 1605 infection (i.p. 2×10^7^ CFU/mouse), numbers of mice (n) are indicated in parentheses. Data were collected from at least three independent experiments, number of biological samples (n) as indicated in parentheses, *, P < 0.05, **, P < 0.01 compared to wild-type. mean ± SEM. Kaplan-Meier estimate was used to compare mice survival rates. Non-parametric t-test was used to compare differences between groups.

**Figure 7.**
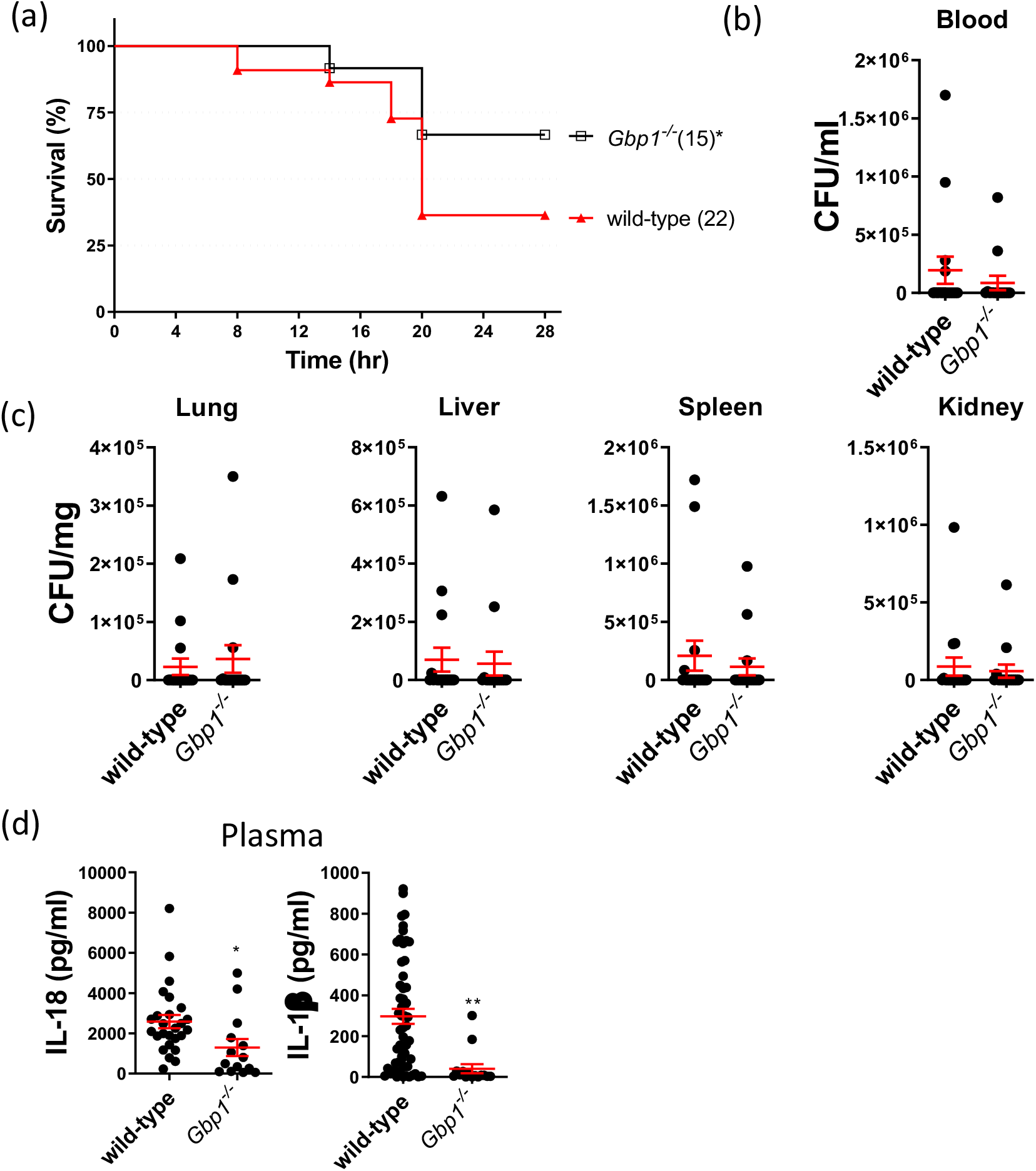
GBP1 drives acute lethality in *A. baumannii*-infected mice. **(a)** Mice survival rate, **(b)** the level of bacteremia, **(c)** bacteria dissemination to different organs, and **(d)** plasma cytokine levels 16-20 hours post *A. baumannii* 1605 infection (i.p. 2×10^7^ CFU/mouse). Data were collected from at least three independent experiments, numbers of mice (n) are indicated in parentheses, *, P < 0.05, **, P < 0.01 compared to wild-type. mean ± SEM. Kaplan-Meier estimate was used to compare mice survival rates. Non-parametric t-test was used to compare differences between groups.

### IFN-inducible guanylate binding protein 1 (GBP1) mediates bacteria killing in Carbapenem and polymyxin resistant bacteria

The IFN-inducible guanylate binding protein - GBP1 - is a cytosolic receptor for LPS and triggers caspase-11-dependent pyroptosis during infection with certain Gram-negative bacteria, such as *Salmonella* Typhimurium ^30 31^ and *Legionella pneumophila* ^32^. GBP1 binds directly to cytoplasmic LPS and promotes an oligomerization state ^33^. Additionally, phosphate groups on the lipid A of LPS play an essential role in promoting GBP1-LPS interaction and activation of the non-canonical inflammasome pathway ^30^. In *A. baumannii* infection, Colistin resistance to antibiotherapy is mediated by LOS via direct binding to lipid A (LpxA) ^21^. We therefore speculated in the context of *A. baumannii* infection that GBP1 would induce caspase-11-dependent pyroptosis and mediate LOS-dependent killing of *A. baumannii*. To determine the role of GBP1 in mediating LOS-dependent killing and caspase-11 inflammasome activation, we first generated *Gbp1*^*-/-*^ knockout strains using CRISPR-Cas9 gene editing technology ^34^. We then assessed survival, bacterial burden, inflammasome activation and pyroptosis. We observed that *Gbp1*^*-/-*^ mice were resistant to *A. baumannii* 1605 infection with a 70% survival rate, no reduction in bacterial burden **(Fig 7a-c)** although had a strong reduction in plasma pro-inflammatory cytokine levels **(Fig 7d)**. Immunoblotting confirmed the activation of caspase-11 was inhibited and GSDMD proteolytic cleavage reduced in *Gbp1*^*-/-*^ BMDMs compared to WT BMDMs **(Fig 8a)**. Additionally, we found a strong reduction in pro-inflammatory cytokine levels at 12 hours post-inoculation **(Fig 8d)**. These data suggest that GBP1 mediates caspase-11 inflammasome activation in response to *A. baumannii* 1605. Next, to assess the role of LpxA in GBP1-mediated killing, we infected WT and *Gbp1*^*-/-*^ BMDMs with an *A. baumannii* LOS-deficient strain carrying a nonsense mutation in *LpxA* (19606 R) and its complement strains (19606R + LpxA or AL 1847, 19606R + V) and *A. baumannii* ATCC 19606 (WT 19606) strain as controls ^21^. We found in BMDMs either 19606R or their complement strains did not induce caspase 1, caspase 11 activation, nor significantly increased GSDMD proteolytic cleavage level or changed their inflammatory responses measured by LDH and pro-inflammatory cytokine levels **(Fig 8a-b, d-f)** from WT 19606 or *A. baumannii* 1605 strains in WT and *Gbp1*^*-/-*^ BMDMs. Overall, it suggests that LpxA is unlikely to be a target for GBP1 bactericidal activity and inflammasome activation. To further confirm this finding, we next assessed direct killing from GBP1 by co-incubating the full-length purified GBP1 protein with *A. baumannii* strains and determined its killing effect. Interestingly, we found the full-length purified mouse GBP1 protein exerted a bactericidal activity *A. baumannii* strains and again no significant change was observed between 19606 WT, 19606R and 19606R+LpxA (**Fig. 8c**). Together, these results suggest that GBP1 exerts a bactericidal activity against *A. baumannii* irrespective of the bacteria’s resistance to carbapenem and polymyxin. Together, our data also show that GBP1 activates caspase-11 inflammasome and also caspase-1 via type I IFN signalling. finally, these experiments indicate that LOS is unlikely to mediate GBP1 bactericidal activity.

**Figure 8.**
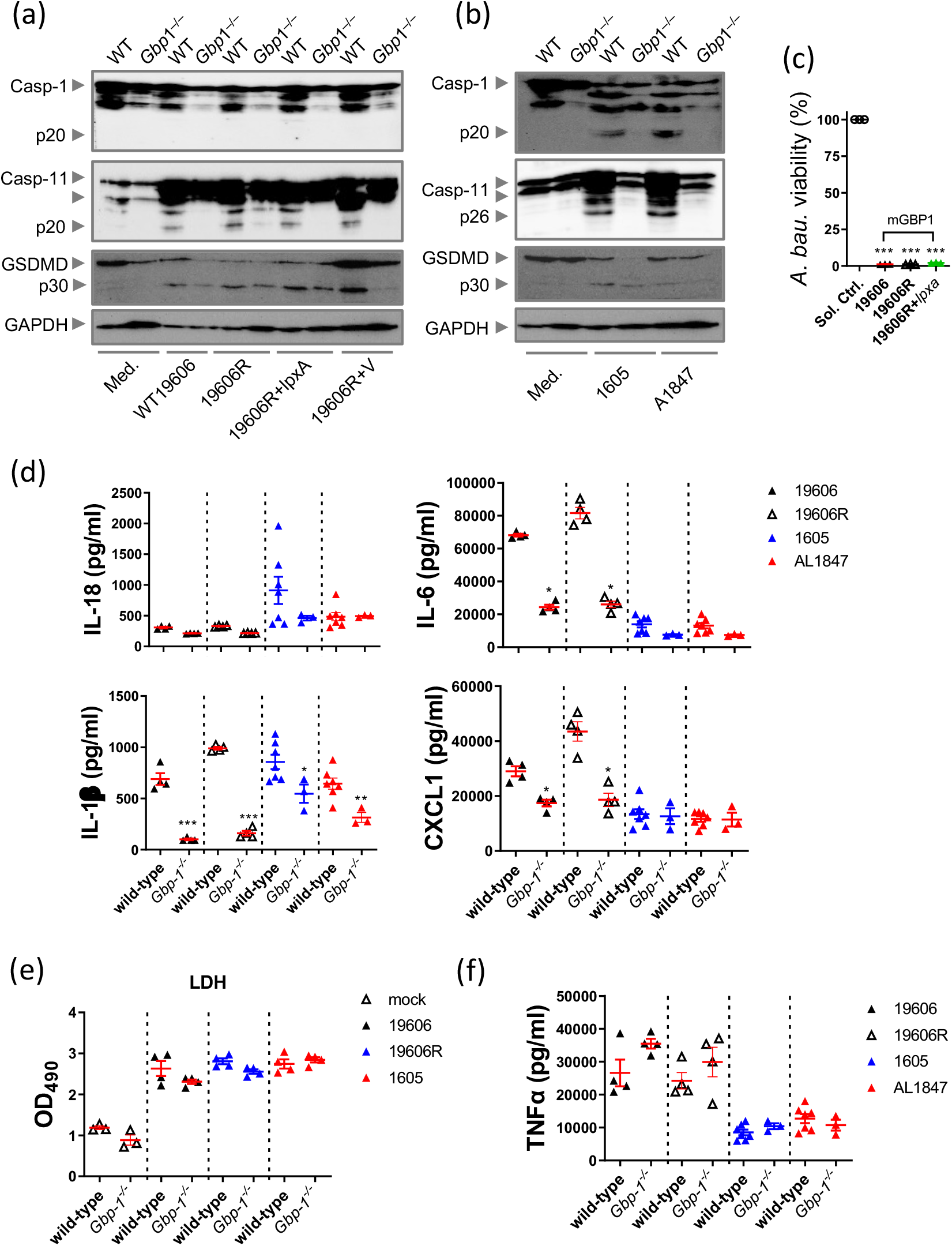
GBP1 drives LPS-independent *A. baumannii* responses. Representative western blots on activated caspases and GSDMD of *A. baumannii* **(a)** 19606, 19606R and 19606R + *LpxA* and **(b)** *A. baumannii* 1605 and A1847 strains at 16 hours post-infection (MOI = 10). (c) *A. baumannii* ATCC 19606 viability post incubation with full-length GBP1 protein, n=3. (d) Cytokine IL-1β, IL-18, IL-6 and CXCL1 levels in supernatants 12 hours post infection (MOI = 10), n = 3-7, each data point represents a replicate. (e) LDH release by mouse BMDM post 12 hours of infection, n=4. (f) TNFα levels post different *A. baumannii* strains infection, n=4. *, P < 0.05, **, P < 0.01, ***, P < 0.001 compared to the respective wild-type. mean ± SEM. Non-parametric t-test was used to compare differences between groups.

## Discussion

*A. baumannii* is a Gram-negative bacterium that causes opportunistic pulmonary and systemic pathologies in humans and is of importance for its resistance to last-resort antibiotics ^3^. Despite the clinical significance of *A. baumannii* in humans, little is known about the role of the innate immune system in host defence against the pathogen. Inflammasome activation is key for innate immune recognition of pathogens and for innate host defences ^35^. While NLRP3 activation has been reported as critical for host immunity in response to *A. baumannii* ^15,36,37^, the role of non-canonical inflammasome activation and cytosolic immunity in sensing *A. baumannii* bacteria remained elusive and poorly understood. A deeper understanding of the host immune defence mechanisms is required to devise potential novel therapies against MDR/XDR bacteria.

Our findings demonstrated that recognition of MDR *A. baumannii* 1605 infection by the host predominantly activates the caspase-11 inflammasome, resulting in NLRP3/ASC inflammasome activation and formation of GSDMD pores, resulting in induction of programmed cell death. Importantly, we found that caspase-11 and IL-1β, via type I IFN, regulate the tight balance between protective and deleterious inflammation. Finally, we discovered that upon bacterial recognition, the cytosolic molecule GBP1 exerts a bactericidal activity against carbapenem-resistant and polymyxin-resistant bacteria (lacking LOS) and to activate the caspase-11 inflammasome. Together our findings have demonstrated that cytosolic innate immune defences mediated by caspase-11, IFN and GBPs are required to facilitate protective inflammation and exert a bactericidal activity against MDR *A. baumannii* bacteria.

Previous reports have identified the requirement of NLRP3/ASC and the caspase-1 canonical pathway ^15,36^ as well as a protective role of caspase-11 in the host inflammatory response to *A. baumannii* ^38^. Our data confirm these findings and establish that NLRP3 inflammasome is exclusively activated by MDR *A. baumannii* 1605. We further investigated the mechanisms of host protection and the role of caspase-11 and GSDMD dependency during MDR *A. baumannii* 1605 infection. In agreement with a previous report ^29^, we found MDR *A. baumannii* 1605 triggers caspase-11 and GSDMD-induced cell death. Our findings and our previous assessment on multiple clinical strains ^39^ however, clearly establishes that the non-canonical inflammasome activation via NLRP3/ASC and GSDMD-caspase-11-mediated cell death are major host defence mechanisms against MDR *A. baumannii*. Intriguingly, we demonstrated IL-1β and not IL-18 is required to regulate a tight balance between protective versus deleterious inflammation, thus controlling severe sepsis. Further, IL-1 was involved in host protection against *Toxoplasma gondii* ^40^ and group B *Streptococcus* infection ^41^ and was reported to play a minor role in the response to drug sensitive *A. baumannii* in a lung pathology model ^36^. To our knowledge, the regulatory role of IL-1β and not IL-18 in controlling beneficial versus deleterious inflammation has not been reported previously for *A. baumannii* infection. Future studies will aim in elucidating the role of IL-1β in MDR and virulent *A. baumannii* infections.

Neutrophils and inflammatory monocytes are abundant resident populations and are rapidly recruited to the infection site to kill micro-organisms ^42^. Previous studies have reported that early recruitment of neutrophils to the lungs was mediated by NLRP3 and was dispensable for the host defence ^43-46^. Our studies revealed that more important than early recruitment to the target tissues, is a balance between effector cell recruitment and their persistence; this balance is responsible for enhanced host resistance against MDR *A. baumannii*. It therefore suggests the innate immune response against *A. baumannii* infection is driven by a tight regulation between effector cell recruitment and programmed cell death mediated by inflammasome activation, controlled by NLRP3/ASC and caspase-11.

Previous work identified IFN is required for host resistance to virulent *A. baumannii* infection by mediating multiple cell death pathways via caspase-11 ^29^. However, the role of cytosolic immunity mediated by IFN molecules for *A. baumannii*-mediated infection is unknown. We did, however, demonstrate an IFN requirement in mediating the cytosolic immune response via caspase-11 activation, cell death and the activation of IFN induced bactericidal proteins in agreement with a previous report ^29^. Importantly, our findings highlight the requirement of cytosolic immunity mediated by GBP1 in the response and bactericidal activity towards MDR *A. baumannii* 1605. Previous work identified human GBP1 as cytosolic receptors for LPS in *S*. Typhimurium and *S. flexeneri* ^30,47,48^ while it was not demonstrated for the mouse GBP1 ^34^. Here, we firstly demonstrated that full-length purified GBP1 exerts a bactericidal activity against LOS-deficient strains resistant to polymyxin, as well as a MDR *A. baumannii* 1605 strain resistant to carbapenems. Our data also suggest that mouse GBP1 is not a receptor for LOS in *A. baumanii* strains, in contrast to previous work demonstrated the direct binding of human GBP1 to LPS ^30,33^. Our finding is in line with previous studies on *Neisseria meningitidis* ^34^ and *Moraxella catarrhalis* LOS and *E. coli* LPS ^49^, demonstrating the lack of direct interaction between mouse GBP1 and the LOS. It therefore suggests that a ligand, other than LOS, directly binds to mouse GBP1 in MDR *A. baumannii* leading to vesicle rupture and caspase-11 activation. Importantly, we demonstrated that GBP1 exerts a bactericidal activity in carbapenem and colistin resistant *A. baumannii*, paving the way to the development of host directed therapy, adjuvant to current antibiotherapy, and perhaps consisting of GBP1 activation using a peptide mimetic or small molecule compounds to enhance MDR/XDR bacteria killing.

In conclusion, we demonstrated a critical role of the caspase-11 non-canonical pathway and cytosolic immunity in mediating bacterial killing, activating the inflammasome and protective pro-inflammatory cytokine production against MDR *A. baumannii* strains resistant to polymyxin and carbapenem.

## Method

### Bacteria strains

*Acinetobacter baumannii* BAA-1605 strain (*A. baumannii* 1605) was obtained from the American Type Culture Collection (ATCC). This strain is a multi-drug resistant to many antibiotics including carbapenems (resistant to ceftazidime, gentamicin, ticarcillin, piperacillin, aztreonam, cefepime, ciprofloxacin, imipenem, and meropemem). This stain was an isolate from sputum of military personnel returning from Afghanistan. *Acinetobacter baumannii* AL 1847, a clinical isolate harbouring a 30bp mutation in *LxpA* resulting in a frameshift mutation, *Acinetobacter baumannii* 19606, AL 1847, an ATCC 19606 derivative harbouring a 30bp mutation in *LxpA* resulting in a frameshift mutation, 19606R (resistant to polymyxin B, carrying a nonsense mutation in LpxA gene), and their complemented strains 19606R-LpxA, 19606R-V (transformed with the empty shuttle vector pWH1266) were obtained from Monash University (Prof John Boyce) and were previously described ^21^. Frozen stocks of bacteria were streaked onto trypticase soy agar plates (Cat. No.: 211768, BD, with 1.5% agar (BD Cat. No.: 281230)) and incubated overnight under aerobic conditions at 37°C. Single colonies were picked and inoculated into 5 ml of trypticase soy broth and were incubated under aerobic conditions at 37°C on an orbital shaker at 250 rpm for 16-18 hours until cloudy. The bacteria were then sub-cultured in 30 ml of trypticase soy broth for further propagation for 4 hours at 37°C, on an orbital shaker at 250 rpm. The bacteria 19606R and 19606R+LpxA were maintained under colistin selection pressure ^21^. Bacterial stocks were prepared by adding 30% of sterile glycerol (Cat. No.: G2025, Sigma-Aldrich), aliquoted into 2 ml vials and frozen at -80°C before use.

### Primary bone marrow derived macrophages (BMDMs)

Mouse bone marrow derived macrophages were used as a mouse macrophage infection model. Total bone marrow was extracted from both mouse femurs, passed through a 70 µm cell strainer (Cat No.:352350, BD Falcon) and centrifuged at 430 relative centrifugal force (rcf) for 5 minutes. The supernatant was discarded. Red blood cells were lysed using 10 ml of 1x red blood cell (RBC) lysis buffer per mouse for 5 minutes during centrifugation.

To promote differentiation into macrophages, bone marrow cells were washed and seeded at 5×10^6^/dish in 10 cm sterile dish in 10 ml of RPMI/10% Foetal Calf Serum (FCS) including 10 ng/ml of mGM-CSF (Cat. No.: 130-095-739, Miltenyi Biotech). Cells were incubated at 37°C 5% CO_2_, 95% relative humidity. The day of isolation is considered as day 0. On day 3, an extra 5 ml of fresh RPMI/10% FCS including 10 ng/ml of mouse GM-CSF was added to replenish the cytokines. On day 6 or 7, adherent cells were collected using RPMI/5 µM EDTA to detach cells from the plates. Cells were seeded in 96-well plates at 1×10^5^/well in RPMI/10% FCS and incubated overnight at 37°C 5% CO_2_, 95% relative humidity, prior to infection.

### Lactic dehydrogenase (LDH) assay

Levels of LDH released by cells were determined using a CytoTox 96 Non-Radioactive Cytotoxicity Assay according to the manufacturer’s instructions (Cat. No.: G1780, Promega). All plates were measured using TECAN Infinite^®^ 200 Pro (Tecan, Männedorf, Switzerland).

### Immunofluorescence

BMDMs were seeded at 4×10^5^/well in sterile 24-well glass bottom plate (Cat. No.: P24-1.5HN, Cellvis) prior to *A. baumannii* inoculation. *A. baumannii* strain 1605 was prepared at multiplicity of infection (m.o.i.) 10 and cells were infected for 24 hours (final volume 1 ml/well) at 37°C 5% CO_2_ in air. Post infection, cells were washed twice with sterile 1xPBS before staining with Zombie Aqua (1:100, 100 μl/sample, Cat. No.: 423101, BioLegend) and Hoechst 33342 (80 micromole (μM)/100 μl/sample, Cat No.: H1399, Invitrogen, Carlsbad, CA, USA) for 30 minutes at room temperature. Post staining, cells were washed twice and fixed in 4% paraformaldehyde (PFA - Cat. No.: 420801, BioLegend, San Diego, CA, USA) for 30 minutes at room temperature. Samples were examined and imaged using a Zeiss Axio Observer with an epifluorescence attachment and a digital camera. Five random fields were taken per well and quantified using Image J with colour deconvolution plugin for mean staining area per channel (ver 1.64r).

### Enzyme-linked immunosorbent assay (ELISA)

Sandwich ELISA was used to measure the release of inflammatory cytokines IL-1β, TNFα and IL-18 in the cell supernatant, cell lysate, or mouse plasma post bacterial infection.

For mouse TNFα (Cat. No.: 88-7324-88, Invitrogen) and IL-1β (Cat. No.: 88-7013-88, Invitrogen), 96-well ELISA plates (Cat. No.: 9018, Corning) were prepared per the manufacturer’s instructions. For the pre-coated IL-18 ELISA (Cat. No.: BMS618-3, Invitrogen), the experiments were performed according to the manufacturer’s instructions.

All plates were measured using TECAN Infinite^®^ 200 Pro (Tecan, Männedorf, Switzerland), with wavelength set at 450 nm.

### RNA extraction and conversion to cDNA

Samples were lysed directly in Trizol reagent (Cat. No.: 15596018, Life Technologies) and stored at -80°C until RNA extraction. RNA extraction was carried out using Qiagen RNeasy kit (Cat. No.: 74134, Qiagen) according to the manufacturer’s instructions. RNA samples were eluted using RNase-free water provided by the kit and then stored at -80°C. RNA was then converted to cDNA following the manufacturer’s instructions for MultiScribe reverse transcriptase (Cat. No.: 4368813, Applied Biosystems). Briefly, RNAse free water was added to 0.5 µg of total RNA to a final volume of 10 µl. 10 µl of reaction mixture containing random primers, dNTPs (dATP, dGTP, dCTP and dTTP) and MultiScribe reverse transcriptase (Cat. No.: 4368813, Applied Biosystems) were added to the RNA solution. The samples were mixed and heated to 25°C for 10 minutes, incubated at 37°C for 2 hours, followed by 85°C for 5 minutes in a thermocycler.

### Real-time reverse transcriptase polymerase chain reaction (real-time RT-PCR)

10 µl of SSOAdvanced™ Universal SYBR^®^ Green Supermix (Cat. No.: 1725275, Bio-Rad), 0.6 µl of 10 µM forward and reverse primer each (Supp. Table 1) and nuclease free water was transferred to each well of a MicroAmp™ fast optical 96-well reaction plate (Cat. No.: 4346907, Applied Biosystems). Diluted cDNAs were added to wells in duplicate while non-template control wells were also loaded. PCR was performed using ABI StepOne™ real-time PCR system, version 2.1 software program (Applied Biosystems, Foster City, CA, USA). Real-time RT-PCR data was analysed using the comparative 2^-∆∆CT^ method^50^ with Gapdh as a housekeeping gene.

### Immunoblotting

Post infection, BMDMs and the collected supernatant sample were lysed in Radioimmunoprecipitation assay buffer (RIPA) lysis buffer supplemented with protease inhibitors, i.e., Complete Protease Inhibitor Cocktail Tablets (Cat No.: 04693132001, Roche) to prevent sample degradation. Samples were boiled with 6x Laemmli buffer containing sodium dodecyl sulfate (SDS) and 100 mM dithiothreitol (DTT) for 5 minutes before storing at -80°C.

Sample was then thawed on ice, and heated to 95°C for 10 minutes after thawing. Each sample was loaded on an individual lane of a 4-15% gradient SDS-PAGE gel (Cat No.: 456-1086, Bio-Rad) in SDS running buffer and run with a constant voltage of 200 volts for approximately 25 minutes until the dye front reached the end of the gel. The resolved proteins in the SDS-PAGE gel were then transferred to a 0.45 µm Polyvinylidene fluoride (PVDF) membrane (Cat No.: 1620115, Bio-Rad) by electroblotting. An electric current of 400 mA was applied to the apparatus for 1.5 hours at 4°C. Following the transfer, the membrane was blocked with 5% (w/v) skim milk in PBS for 1 hour at room temperature to prevent non-specific binding of Immunoglobulins (Ig).

The PVDF membrane was incubated with primary mouse anti-mouse caspase-1 (1:1000, Cat. No.: 106-42020, Adipogen), caspase-11 (1:1000, Cat. No.: NB120-10454, Novusbio), or Glyceraldehyde 3-phosphate dehydrogenase (GADPH) (1:1000, Cat No.: MAB374, Merck Millipore), GSDMD (1:3000, Cat No.: ab209845, Abcam), diluted in 1% (w/v) skim milk in PBST (PBS with 1% Tween-20) overnight at 4°C, with gently rocking. PVDF membranes were then incubated with horseradish peroxidase-conjugated secondary antibody (1:5000) for 1 hour at room temperature. Immunoreactive proteins were detected by applying ECL Western blotting Detection Reagent (Cat No.: 1705060, Bio-Rad) or SuperSignal™ West Femto Maximum Sensitivity Substrate (Cat. No.: 34096, Thermo Fisher Scientific). The ChemiDoc™Touch Imaging System (BioRad) was used for all blots.

### Recombinant protein expression and purification

The BL21(DE3) *E. coli* strain (C2527H, NEB) was transformed with pET-28a(+)-TEV plasmid containing the sequence for mouse GBP1 (mGBP1) and transformants were selected with 50 µg/ml kanamycin (10106801001, Roche). A single colony was used to inoculate a starter culture of 10 ml LB_Kan_ broth (LB broth + 50 μg/ml kanamycin) which was incubated at 37°C, shaking (180 rpm) overnight. The overnight culture was diluted 1:100 into 800 ml of LB_Kan_ broth and incubated at 37°C, shaking (180 rpm) for 2-3 hours until an OD_600_ of 0.7 was obtained. Cultures were cooled to room temperature, expression was induced by adding isopropyl β-D-1-thiogalactopyranoside (0.5 mM; IPTG, Roche) and the incubation continued at 18°C with shaking (180 rpm) overnight. The culture was centrifuged (5000 × *g*, 20 minutes, 4 °C) to pellet the bacteria and stored at −80°C until required. The cell pellet was resuspended in lysis buffer (50 mM NaH_2_PO_4_, 300 mM NaCl, 10 mM imidazole, 5% glycerol (v/v), 5 mM MgCl_2_, 0.01% Triton X-100, pH 8.0) supplemented with lysozyme (250 µg/ml), Benzonase nuclease (50 U/ml) and protease inhibitor cocktail (11697498001, Roche) and incubated with gentle agitation at 4°C for 1 hour. Cells were subsequently disrupted by sonication and centrifuged (18,000 × *g*, 30 minutes, 4°C) to pellet cellular debris. The supernatant was passed through a 0.22 µm filter (SLGP033RS, Merck) and mGBP1 was purified using Ni-NTA agarose resin (30210, Qiagen) as per the manufacturers’ instructions. The purity of eluted proteins was analyzed by SDS-PAGE and Coomassie blue staining. Purified proteins were dialyzed in DPBS (14190, ThermoFisher) containing 20 mM Tris and 20% glycerol (v/v), pH 7.5.

### Antimicrobial assays

For bacterial viability assays, overnight cultures of *A. baumannii* were washed and resuspended with PBS to a concentration of 1×10^6^ CFU/ml. Bacteria were then treated with solvent control (PBS) or GBP1 at 300 µg/ml and incubated at 37°C for 6 hours. Treated bacteria were serially diluted, plated onto trypticase soy agar plates, and incubated overnight at 37°C. Colonies were enumerated the following day.

### Mice

C57BL/6 mice and *Gsdmd*^*-/-*^ mice carrying a missense mutation impairing pore formation but not proteolytic cleavage ^51^ were sourced from The Australian National University. *Nlrp3*^*-/-*52^, *Casp1*^*-/-*53^, *Casp1/11*^*-/* 53^, *Casp11*^*-/-*54^, *Il18*^*-/-*27^, *Il1r1*^*-/-*28^ and *Ifnar*^*-/-* 55^ mice were sourced from The Jackson Laboratory. *Il1r/IL18*^-/-^ were generated via crossing *Il18*^*-/-*^ and *Il1r1*^*-/-*^ mice. *Asc*^*-/-*9^ mice were sourced from the University of Queensland. *Gbp1*^-/-^ was generated by CRISPR-Cas9 gene editing technology and was previously described ^34^.

All mice are on, or backcrossed to, the C57BL/6 background for at least 10 generations. Male and female mice of 8-12-weeks old were used. Mice were bred and maintained at The Australian National University under specific pathogen-free conditions. All animal studies were performed in accordance with the National Health and Medical Research Council code for the care and use of animals under the Protocol Number A2018-08 and A2021-14 approved by The Australian National University Animal Experimentation Ethics Committee.

### *In vivo* infection

*A. baumannii* was streaked onto trypticase soy agar plates and incubated at 37°C overnight for isolation of single colony. Single colonies of *A. baumannii* were picked and inoculated into 5 ml of trypticase soy broth and incubated at 37°C 16-18 hours on shaker at 220 rpm for bacterial propagation. 5 ml of the bacterial broth was diluted 1:5 with fresh trypticase soy broth the next day and incubated for further 2 hours to ensure most of the bacteria cells are in log phase of growth. After incubation, bacteria were collected and wash with sterile 1xPBS at 2,800 rcf for 30 minutes. Mice were infected via intraperitoneal injection of *A. baumannii* (200 µl/mouse, 2×10^7^ CFU/mouse). The mice were monitored every 4 hours until 28 hours post infection. Observations consistent with illness during monitoring include coat condition (ruffles), hydration levels (whether the mice were eating or drinking), and activity level (whether the mice are moving, i.e. if there’s slowing in movement). Each of these categories was scored independently between 0 (normal) and 2 (very ill).

Mice were humanely euthanised when they were considered a score of 2 for any of these categories. Approximately 20-50 mg of organs (20-50 mg of liver, one lung, the spleen and one kidney) were isolated, weighed and filtered through a 70 μm nylon mesh cell strainer in 1 ml of sterile 1xPBS before serial dilutions in 1xPBS and enumeration on trypicase soy agar plates.

### Characterisation of effector cell populations in mice lungs post *A. baumannii* infection

One lung was excised from an infected mouse and rinsed with sterile 1xPBS. The lung was then finely diced and placed in 1 ml of 1 mg/ml Collagenase P/RPMI (Cat. No.: 11-213-873-001, Roche) and incubated at 37°C for 30 minutes. The digested lung was mashed using a 3 ml syringe plunger through a 70 μm nylon mesh cell strainer. The resultant cell suspension was centrifuged at 300 rcf for 5 minutes. The cell pellet was resuspended in 0.5 ml of 1x RBC lysis buffer and centrifuge at 300 rcf for 5 minutes. Lung cells were transferred to a 1.5 ml microfuge tube for cell surface marker staining.

For intracellular viability staining, 100 μl of Zombie Aqua (1:1000, Cat. No.: 423101, BioLegend, San Diego, CA, USA, 1:1000) was added to each sample for 10 minutes at room temperature. Cells were washed with 1xPBS before proceeding with cell surface marker staining. The cells were treated with Fc block (Cat. No.: 553142, BD Pharmingen) (5 µl/sample) for 10 minutes on ice. After incubation, Fc block was removed, and a cocktail of conjugated primary antibodies Ly6g-APC-Cy7 (1:500, Cat. No: 127624, BioLegend), CD11b-PE-Texas Red (1:500, Cat. No: 101256, Biolegend), Ly6c-Alexa Flour 405 (1:500, Cat. No: 48-5932-82, Thermo), was added directly into relevant samples for 30 minutes on ice in the dark. The cells were washed with MTRC buffer and centrifuged for 5 minutes, prior to fixation with 4% PFA for 30 minutes at room temperature in the dark. The total cell population was collected on FACS Fortessa platform (Becton Dickinson). The analysis was performed using FlowJo software (ver. 10.8.1). The cell population was gated using forward scatter and side scatter to exclude debris, followed by doublet exclusion to characterise single cells (**Supp. Fig. 1a**). This population was then gated for different neutrophil populations (**Supp. Fig. 1c and 1d**) and inflammatory monocytes (**Supp. Fig. 1b and 1d**) according to the cell surface marker expression. When characterising cell death using Zombie aqua, cells were further gated based on Zombie aqua fluorescence (**Supp. Fig. 1b and 1c)**.

### Statistical analyses

The GraphPad Prism 8.0 software was used for data analyses. Data are shown as the mean ± SEM. Statistical significance was determined by t-tests (two-tailed) for two groups or one-way ANOVA (with Dunnett’s or Tukey’s multiple comparisons tests) for three or more groups. Survival curves were compared using the log-rank test. A p-value<0.05 was considered statistically significant.

## Acknowledgements

The authors would like to thank Prof John Boyce and Mrs Amy Wright (Monash University, Australia) for providing the strains 19606, 19606R, AL 1847 and 19606+LpxA. The authors thanks Dr Harpreet Vora, Mr Michael Devoy from the flow cytometry at JCSMR and the Australian Phenomics facility for technical assistance. FJ.L. was supported by an ANU-Taiwan scholarship.

## Author contribution

F-J.L., L.S. and G.B. conceived the study. F-J.L., L.S., A.M. and D.E.T. performed the experiments. F-J.L., L.S., S.M.M. and G.B. conducted the analysis. S.M.M provided mice and cells. F-J.L. and G.B. wrote the manuscript. G.B provided the overall supervision of the work. All authors provided feedback and approved the manuscript.

## Competing interests

The authors declare no competing interests.

## Supplementary Tables

**Supplementary Table 1.**
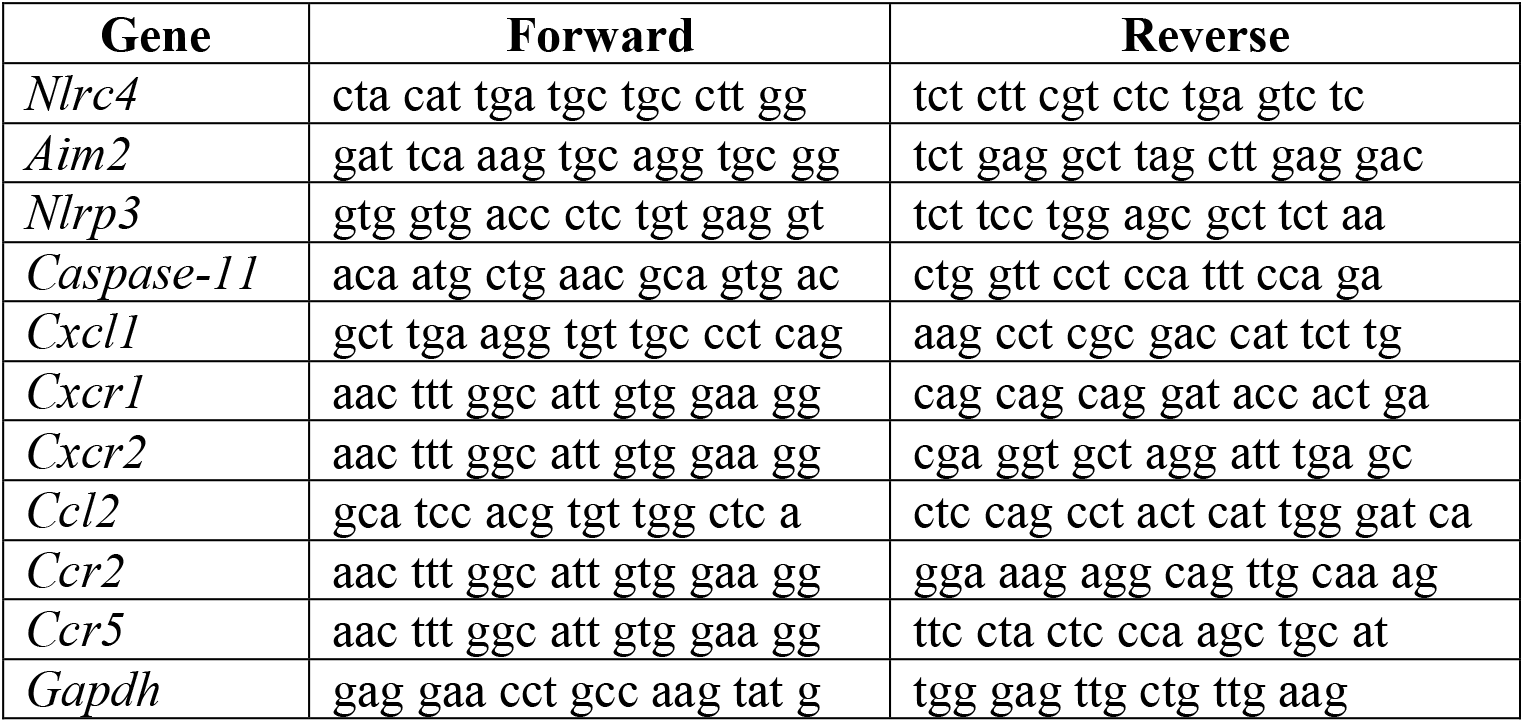
qPCR primers used in this study

## Supplementary Figures

**Supp. Figure 1.**
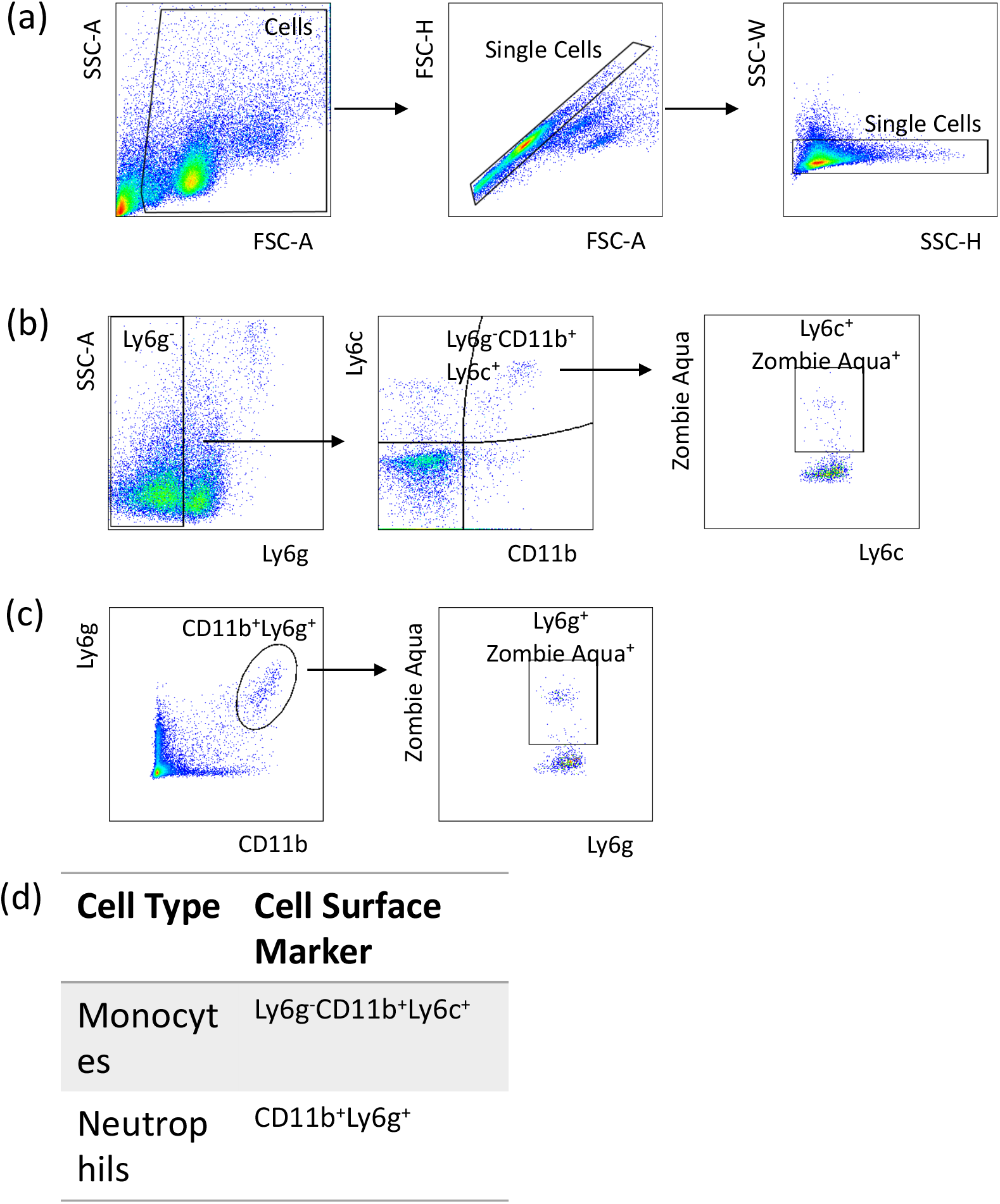
Gating strategy for defining effector cell subpopulations by flow cytometry. Briefly, doublets were excluded using Forward Scatter (FSC) and Side Scatter (SSC) as shown in **(a)** before gating for specific effector populations based on CD11b versus **(b)** Ly6c or **(c)** Ly6g expression.

**Suppl. Figure 2.**
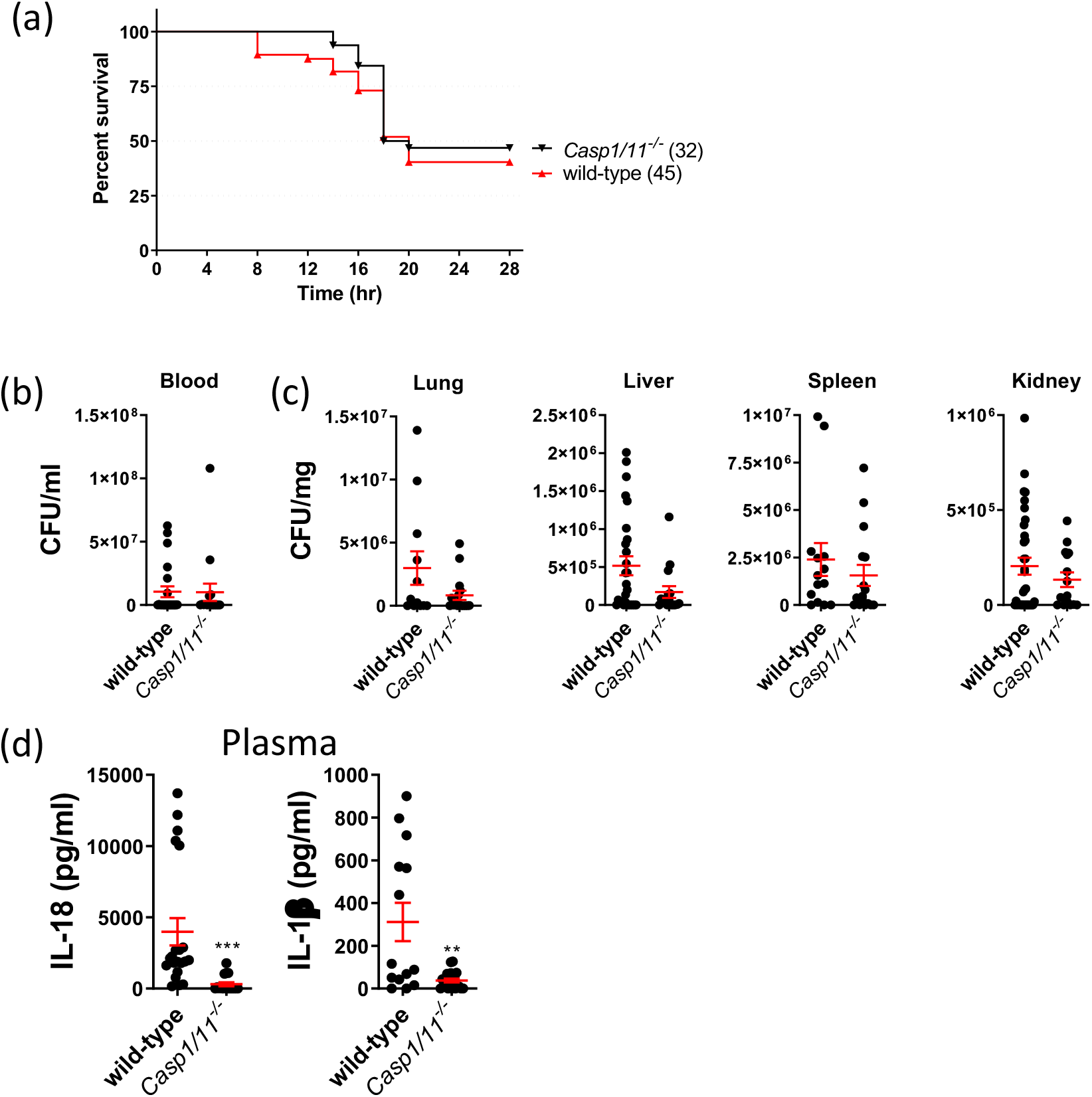
Caspase1/1 mice are equally susceptible to *A. baumannii* infection. **(a)** *Caspase1/11*^-/-^ mice survival rate, **(b)** the level of bacteremia, **(c)** bacteria dissemination to different organ, and **(d)** plasma cytokine levels 16-20 hours post *A. baumannii* 1605 infection (i.p. 2×10^7^ CFU/mouse). Data were collected from at least three independent experiments, n as indicated in parentheses, **, P < 0.01, ***, P < 0.001 compared to wild-type. mean ± SEM. Kaplan-Meier estimate was used to compare mice survival rates. Non-parametric t-test was used to compare differences between groups.

**Suppl. Figure 3.**
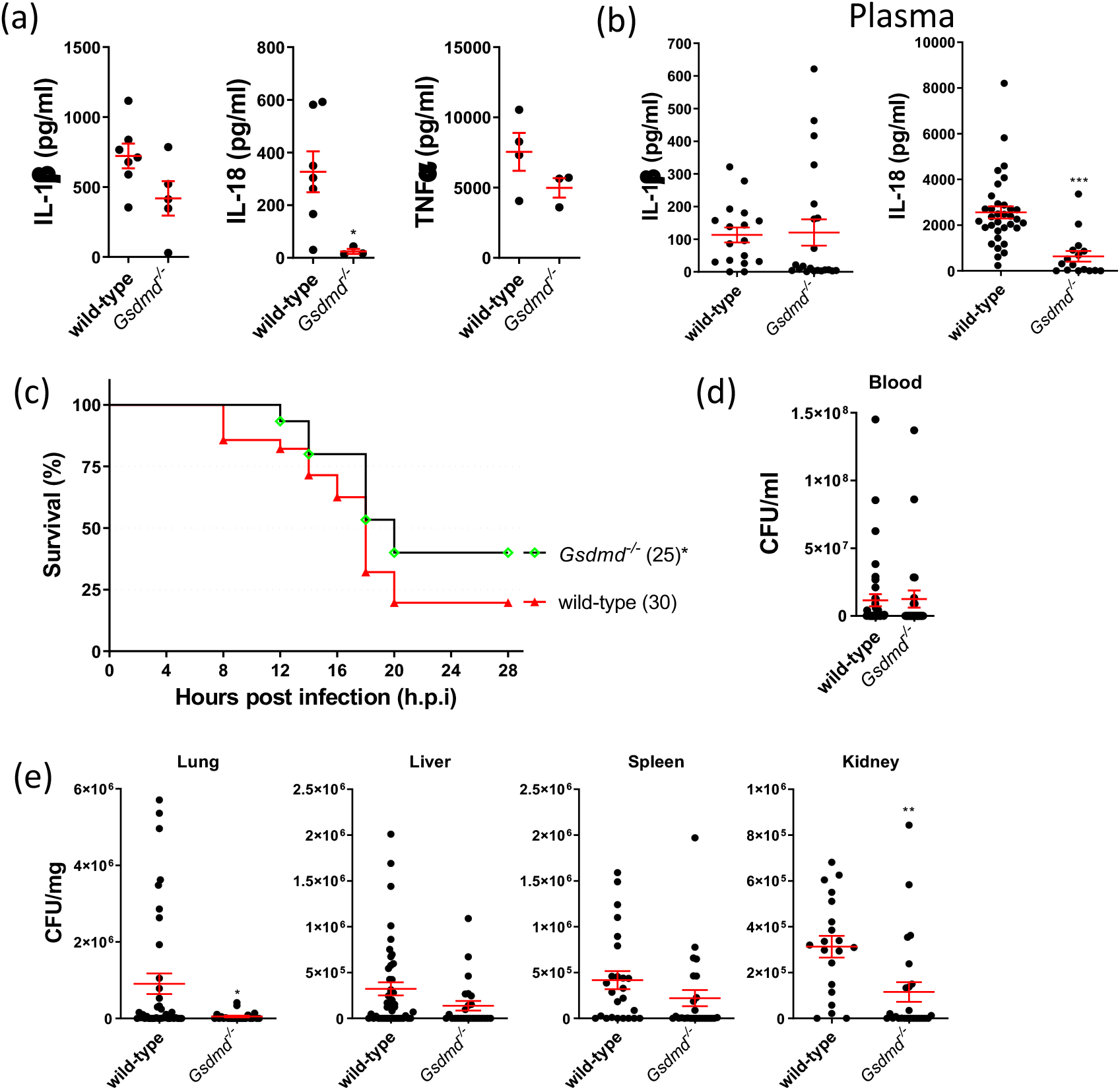
Deleterious inflammation drives acute lethality in *A. baumannii*-infected mice. **(a)** BMDM cytokine levels IL-1β, IL-18 and TNFα in supernatants post 12 hours *A. baumannii* infection (m.o.i.=10), n = 5. **(b)** Mouse plasma cytokine levels, **(c)** survival rate, **(d)** the level of bacteremia and (e) bacteria dissemination to different organs 16-20 hours post *A. baumannii* 1605 infection (i.p. 2×10^7^ CFU/mouse). Data were collected from at least three independent experiments, n as indicated in parentheses, *, P < 0.05, **, P < 0.01, ***, P < 0.001 compared to wild-type. mean ± SEM. Kaplan-Meier estimate was used to compare mice survival rates. Non-parametric t-test was used to compare differences between groups.

**Supp. Figure 4.**
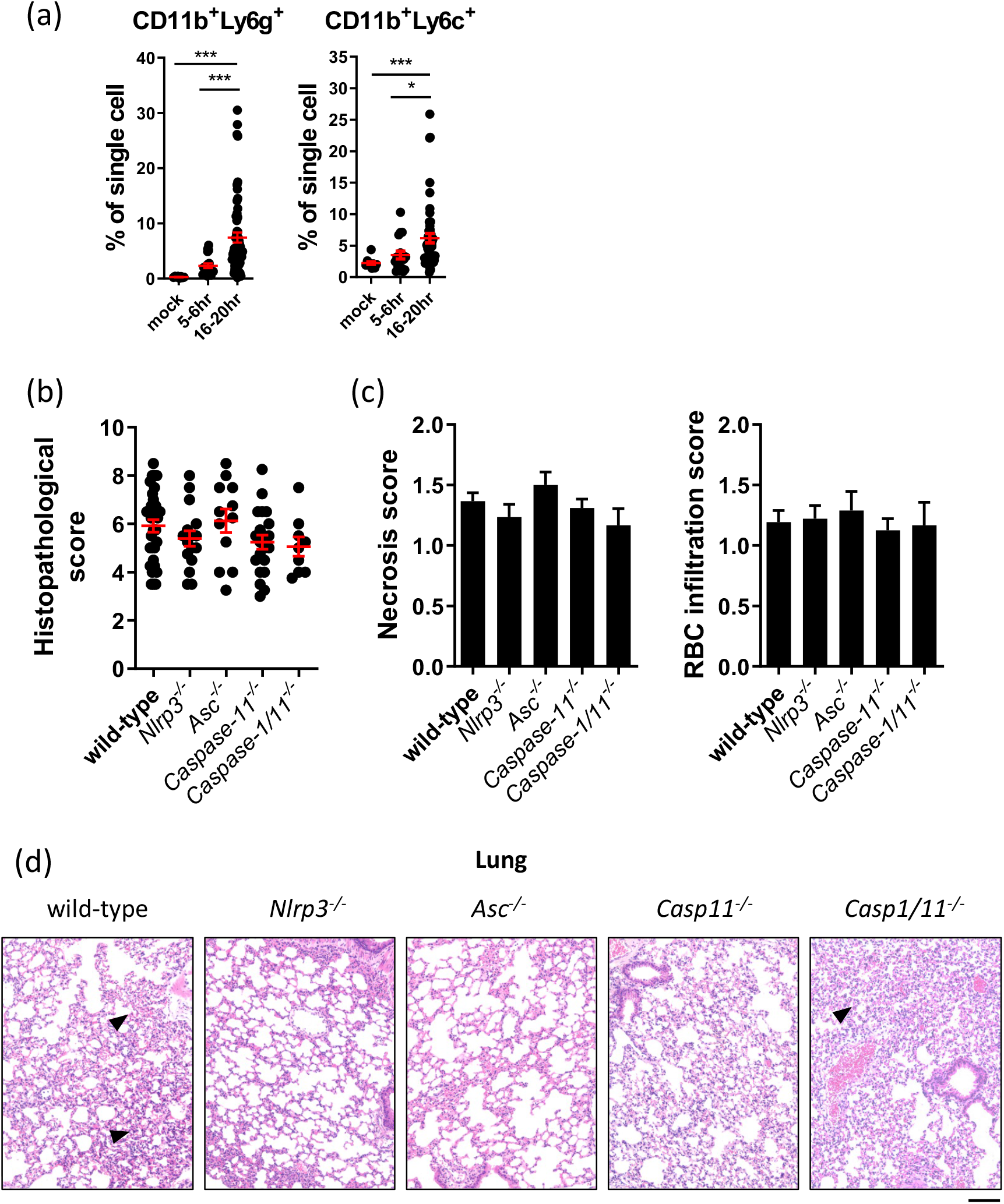
Recruitment of effector cells does not contribute to lung lesions. **(a)** Recruitment of neutrophils and inflammatory monocytes to the lung, **(b)** Lung histological scores, **(c)** necrosis or red blood cell (RBC) infiltration score and the **(d)** representative H&E staining of infected C57BL/6 mice 16-20 hours post *A. baumannii* 1605 infection (i.p. 2×10^7^ CFU/mouse). Arrowhead: immune cell infiltrate. n10-20 n=6-12, each data point represents a replicate. For each group, scale bar: 20 µm. Non-parametric t-test was used to compare differences between groups.

**Supp. Figure 5.**
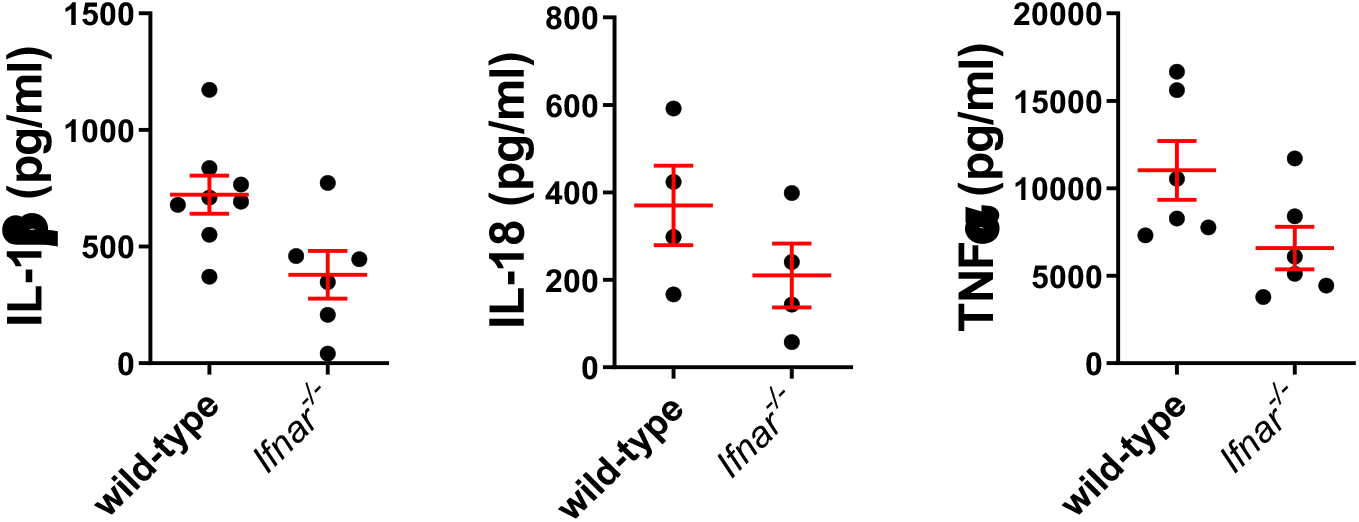
Absence of type I IFN signalling does not alter cytokine levels. Levels of cytokines IL-1β, IL-18 and TNFα in BMDM supernatants 12 hours post *A. baumannii* infection (m.o.i. 10), n = 4-8, each data point represents a replicate, mean ± SEM.

**Supp. Figure 6.**
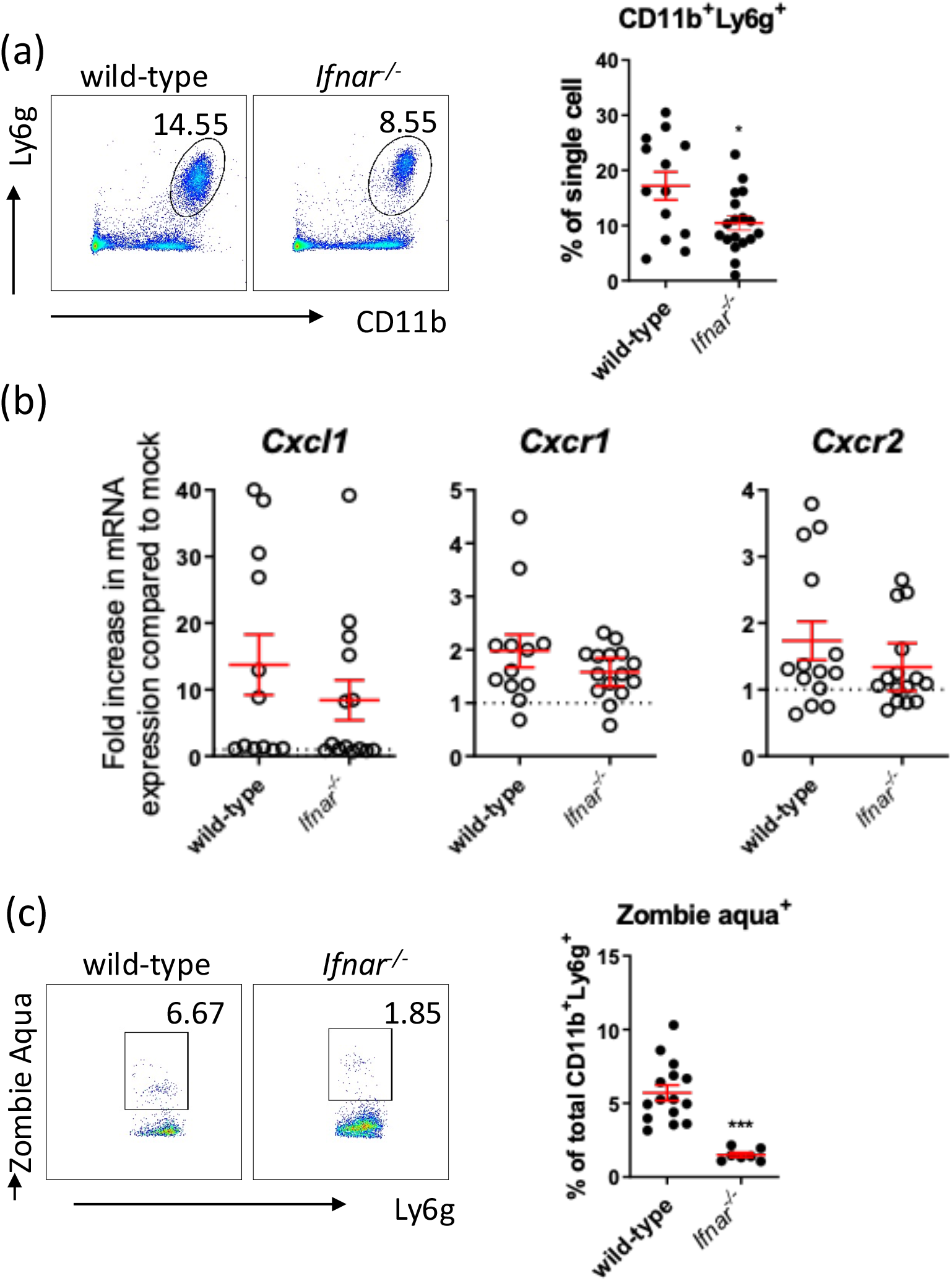
Absence of type I IFN signalling decreases neutrophil recruitment and neutrophil death. Flow cytometry quantification of **(a)** neutrophils (CD11b^+^Ly6g^+^) **(b**) qPCR quantification of induction of neutrophil chemokine and chemokine receptors **(c)** neutrophil cell death, in mice lung 14-20 hours post *A. baumannii* 1605 infection (i.p. 2×10^7^ CFU/mouse). Data were collected from at least three independent experiments. *, P < .05, ***, P < .001, mean ± SEM, n = 10-17, each data point represents a replicate. Non-parametric t-test was used to compare differences between groups.

**Supp. Figure 7.**
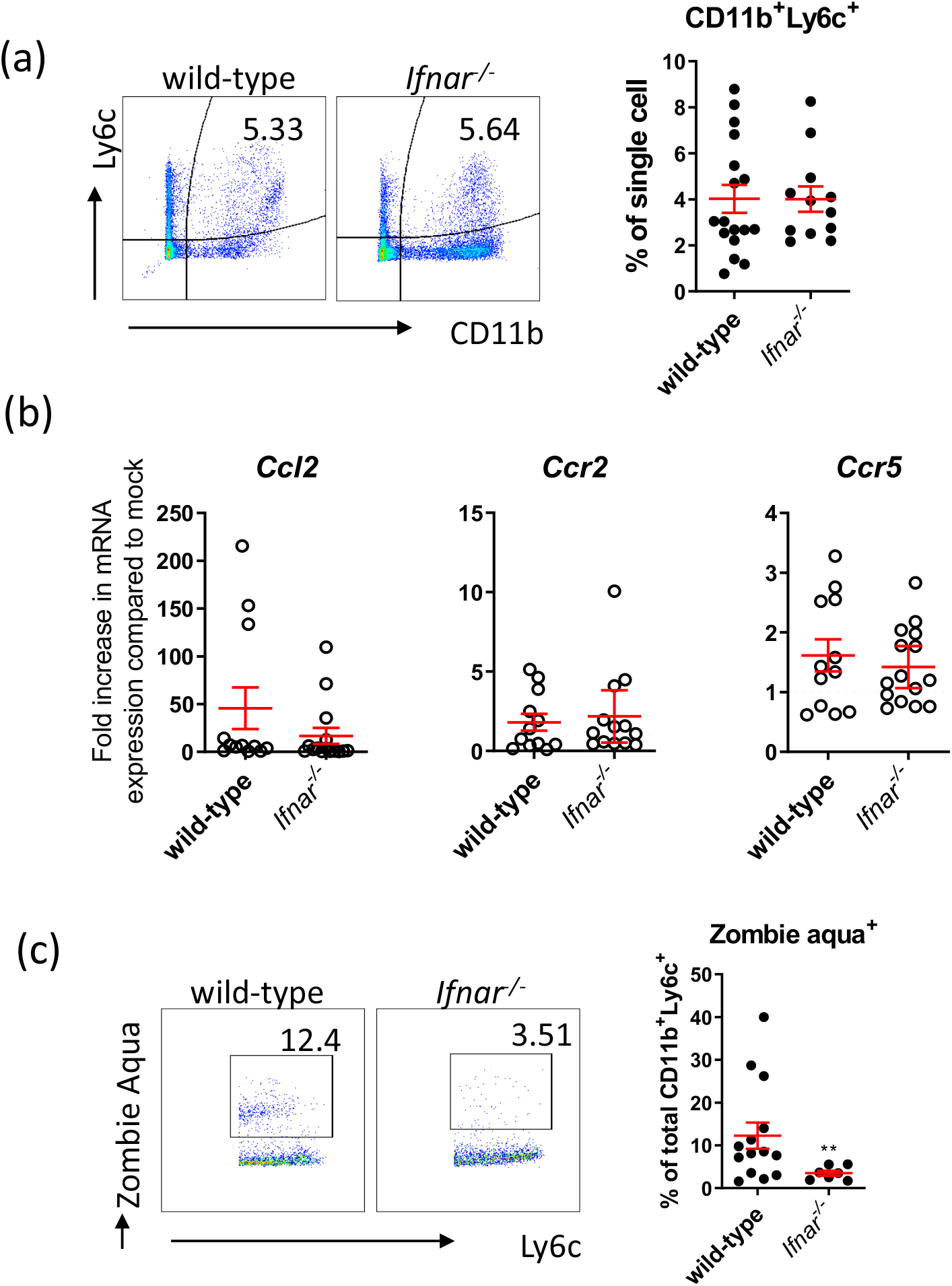
Absence of inflammasome signalling decreases inflammatory monocyte death only. Flow cytometry quantification of **(a)** inflammatory monocytes (CD11b^+^Ly6c^+^) **(b)** qPCR quantification of induction of inflammatory monocytes chemokine and chemokine receptors. **(c)** Inflammatory monocytes cell death, in mice lung post 14-20 hours of *A. baumannii* 1605 infection (i.p. 2×10^7^ CFU/mouse). Data were collected from at least three independent experiments, n = 12-17 for each group, n=6-12, each data point represents a replicate. **, P < 0.01, mean ± SEM. Non-parametric t-test was used to compare differences between groups.

**Supp. Figure 8.**
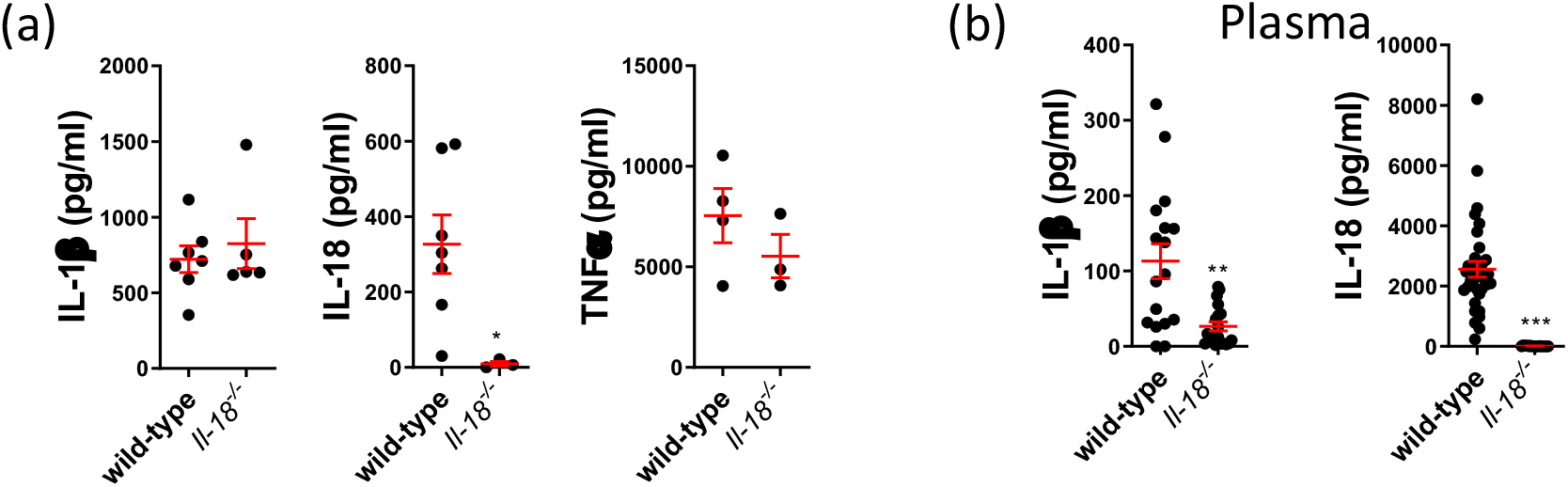
IL-18-knockout mice showed reduced IL-1β. **(a)** Cytokine levels secreted by BMDM into culture supernatant post 12 hour of *A. baumannii* infection (m.o.i = 10), n=4-7, each data point represents a replicate. **(b)** Plasma pro-inflammatory cytokine levels post 16-20 hours of *A. baumannii* inoculation (i.p. 2×10^7^CFU/mouse), n = 20. *, P < .05, ***, P < .001, mean ± SEM. Non-parametric t-test was used to compare difference between group. Non-parametric t-test was used to compare difference between group.

## Notes

### Competing Interest Statement

The authors have declared no competing interest.

